# Parallel Distributed Networks Resolved at High Resolution Reveal Close Juxtaposition of Distinct Regions

**DOI:** 10.1101/475806

**Authors:** Rodrigo M. Braga, Koene R. A. Van Dijk, Jonathan R. Polimeni, Mark C. Eldaief, Randy L. Buckner

## Abstract

Examination of large-scale distributed networks within the individual reveals details of cortical network organization that are absent in group-averaged studies. One recent discovery is that a distributed transmodal network, often referred to as the ‘default network’, is comprised of two separate but closely interdigitated networks, only one of which is coupled to posterior parahippocampal cortex. Not all studies of individuals have identified the same networks and questions remain about the degree to which the two networks are separate, particularly within regions hypothesized to be interconnected hubs. Here we replicate the observation of network separation across analytical (seed-based connectivity and parcellation) and data projection (volume and surface) methods in 2 individuals each scanned 31 times. Additionally, 3 individuals were examined with high-resolution fMRI to gain further insight into the anatomical details. The two networks were identified with separate regions localized to adjacent portions of the cortical ribbon, sometimes inside the same sulcus. Midline regions previously implicated as hubs revealed near complete spatial separation of the two networks, displaying a complex spatial topography in the posterior cingulate and precuneus. The network coupled to parahippocampal cortex also revealed a separate region directly within the hippocampus at or near the subiculum. These collective results support that the default network is composed of at least two spatially juxtaposed networks. Fine spatial details and juxtapositions of the two networks can be identified within individuals at high resolution, providing insight into the network organization of association cortex and placing further constraints on interpretation of group-averaged neuroimaging data.

---

The association cortex is comprised of distributed networks that are spatially organized as a set of parallel networks involving temporal, parietal, frontal, and anterior and posterior midline regions (Yeo et al. 2011; Power et al. 2011; Doucet et al. 2011; Margulies et al. 2016; see also Goldman-Rakic 1988). Recently, repeated scanning within individuals has delineated networks with high spatial precision (Laumann et al. 2015; Braga and Buckner 2017; Gordon et al. 2017; see also Fedorenko et al. 2010; 2012; Peer et al. 2015; Michalka et al. 2015; Huth et al. 2016). Using such procedures, we discovered that a distributed transmodal network, often called the ‘default network’ (DN; Buckner et al. 2008;

Andrews-Hanna et al. 2014; Raichle 2015), is likely comprised of at least two parallel networks (Braga and Buckner 2017). The two networks are closely juxtaposed throughout the cortex and interdigitated in a way that suggests why features of the more detailed organization are lost in group-averaged data. The two networks were referred to as Network A and Network B. One important distinction between the networks is that Network A is coupled with the posterior parahippocampal cortex (PHC), while Network B is not, suggesting that they may be functionally specialized.

Since our observation of parallel interdigitated networks, multiple studies of individuals have revealed related distinctions but none with the same parallel structure distributed throughout the cortex (e.g., see the ‘Default A’ and ‘Default B’ networks in Fig. 4 of Kong et al. 2018; the ‘Default’ network in Fig. 3 of Gordon et al. 2017; also Wang et al. 2015; Laumann et al. 2015). Thus the details of network organization require replication and further exploration. Of particular interest are zones of cortex than have been variably identified as ‘cores’ or ‘hubs’ of the DN. The designations reflect results that suggest the anterior and posterior midline are convergence zones of functional coupling between distributed systems (e.g., Buckner et al. 2009; Tomasi and Volkow 2010; see also Braga et al. 2013) and are active across multiple task classes (e.g., Andrews-Hanna et al. 2010; 2014) that otherwise show specificity (see also Power et al. 2013). The recent reevaluation of the functional anatomy within individuals necessitates a more detailed look at the anterior and posterior midline with higher spatial resolution methods that are able to disambiguate whether separate networks converge on common regions or alternatively whether separate functional regions lay side-by-side (Fedorenko et al. 2012; Michalka et al. 2015).

High resolution imaging at high field offers a potential means to refine our understanding of network organization, in particular for regions that have complex organization including the anterior and posterior midline. Two prior studies have examined functional connectivity of the DN at high field. Hale et al. (2010) demonstrated that the DN could be identified at 7T with high signal-to-noise, requiring lower spatial smoothing than conventional 3T imaging. Their proof-of-concept analysis focused on similarities between 3T group data and 7T. While robustly identifying the canonical DN, their analysis did not reveal any indications of substructure within the DN (which was not the focus of the study). De Martino et al. (2001) extensively analyzed the spatial properties of the DN within individuals at 7T and made two relevant observations. First, the DN could be resolved within individuals using high-resolution acquisition (1-1.5mm) and minimal spatial smoothing. Second, they found that the DN regions were largely located within the gray matter often following the cortical ribbon. Their analyses focused on the DN as defined in group-averaged data and therefore they did not seek or detect spatially juxtaposed networks within the canonical DN. Nonetheless, the observation that the regions followed the cortical gray matter ribbon bodes well for the use of high field fMRI to resolve spatial details of functional connectivity networks within individuals. Of particular interest, Peer et al. (2015) examined task-based responses in or near regions associated with the DN at high resolution. They observed a complex organization that included both domain-general as well as domain-specific regions along the posterior midline

In the present study, we acquired new data in order to explore further the hypothesis that there exist parallel interdigitated networks within the canonical group-defined DN. Critically, in addition to 3T fMRI data that could replicate our earlier analyses, we also explored high-resolution fMRI data acquired over multiple runs at 7T to delineate the parallel networks in relation to the topography of the cortical ribbon itself. Closely juxtaposed and often minimally overlapping portions of the two networks were identified along the cortical ribbon, with neighboring regions sometimes resolved inside the same sulcus.

## Methods

### Participants

Five healthy right-handed adult women were recruited from the greater Boston community and screened to exclude a history of neurological and psychiatric illness. Data were collected as part of two separate studies. The 3T study acquired fMRI data using similar procedures to Braga and Buckner (2017). The 7T study acquired data at higher spatial resolution. Participants provided written informed consent using procedures approved by the Institutional Review Board of Harvard University (3T study) and Partners Healthcare (7T study). 3T study participants were each scanned across 31 separate MRI sessions (n=2; ages 22 and 23; over 28 weeks for one individual and 40 weeks for the other). 7T study participants were each scanned in a single MRI session (n=3; ages 19–28).

### 3T MRI Data Acquisition

Data were acquired at the Harvard Center for Brain Science on a Siemens Prisma-fit 3T MRI scanner using the vendor’s 64-channel phased-array head-neck coil (Siemens, Erlangen, Germany). The vendor’s head-coil-shaped pillow was used with additional eggshell foam padding on the top and side of the head for immobilization. The scanner room was illuminated to enhance alertness. Eyes were monitored and video-recorded using an Eyelink 1000 Core Plus with Long-Range Mount (SR Research, Ottawa, Ontario, Canada). A 4-point scale was used to record participant’s level of arousal during each run based on the frequency and duration of eye closures. A video of the eye tracker output (showing eye closures) was also retained in order to assess and quantify compliance. Functional runs were flagged for exclusion if 1) maximum absolute motion exceeded 2mm, 2) slice-based temporal signal to noise ratio was less than or equal to 135, and3) the value for maximum absolute motion or signal-to-noise ratio represented an outlier when values from all runs were plotted together. The raw data from flagged runs were then visually checked for motion artifacts and excluded if these were deemed to be severe. Following this procedure, no runs were excluded for S1 and 1 run was excluded for S2. Due to data loss (failures) during processing, 3 additional runs were discarded for S1 and 9 additional runs were discarded for S2, leaving a total of 59 and 53 runs, respectively.

Blood oxygenation level-dependent (BOLD) fMRI (Kwong et al. 1992; Ogawa et al. 1992) data were acquired using a multi-band gradient-echo echo-planar pulse sequence (Setsompop et al. 2012): TR 1000 ms, TE 32.6 ms, flip-angle 64°, 2.4 mm isotropic voxels, matrix 88 × 88 × 65, multi-slice 5× acceleration. The custom sequence was generously provided by the Center for Magnetic Resonance Research (CMRR) at the University of Minnesota. Minimization of signal dropout was achieved by automatically (van der Kouwe et al. 2005) selecting a slice plane 25° from the anterior-posterior commissural plane towards the coronal plane (Weiskopf et al. 2006; Mennes et al. 2014). A rapid T1-weighted anatomical scan was acquired in each session using a multi-echo MPRAGE three-dimensional sequence (van der Kouwe et al. 2008): TR 2200 ms, TE 1.57, 3.39, 5.21,7.03 ms, TI 1100ms, flip angle 7°, 1.2 mm isotropic voxels, matrix 192 × 192 × 144, in-plane GRAPPA acceleration 4. A dual-gradient-echo B0 fieldmap was acquired to correct for sus-ceptibility-induced gradient inhomogeneities: TE 4.45, 6.91 ms with matched slice prescription/spatial resolution to the BOLD sequence.

Each subject’s 31 MRI sessions included at least two BOLD runs of fixation each lasting 7m 2s. Participants were instructed to remain still, stay awake and to maintain fixation on a centrally presented black crosshair viewed on light gray background through a mirror attached to the head coil. The screen was adjusted to ensure comfortable viewing. Sessions included additional task runs (‘n-back’ working memory and visuomotor tasks) and an arterial spin labeling sequence that were not analyzed here.

The broader scope of the 3T data (which extended beyond the present aims) was to assess the effects of transcranial magnetic stimulation (TMS) on functional connectivity in individuals (stimulation administered before the MRI sessions). The effects of TMS are minimal and there has been no detectable effect on topography of cortical organization, so the data were treated as standard rest fixation for the present purposes (confirming this practical decision, the present findings do not differ if only data after sham stimulation are examined). For seed-based functional connectivity analyses in both the volume and surface, the 3T data were divided into four independent datasets (n = 12 in each dataset, yielding 1h 22m of data after exclusion of the initial 12 volumes in each run for T1 equilibration effects). These multiple datasets were used to replicate the observed network distinctions. Best estimate seed-based functional connectivity maps were created by averaging the maps produced from the four datasets in each subject. For *k*-means clustering in both the volume and surface, 59 runs were simultaneously included for S1 (yielding 6h 43m 10s of usable data) and 53 runs for S2 (6h 02m 10s of usable data).

### 7T MRI Data Acquisition

Data were acquired at the Athinoula A. Martinos Center for Biomedical Imaging on a whole-body 7T MRI scanner (Siemens, Erlangen, Germany) equipped with SC72 body gradients (70 T/m maximum gradient strength and 200 T/m/s maximum slewrate) and a 32-channel RF loop coil head array (Keil et al. 2010) for reception and detunable band-pass birdcage coil for transmission. A pillow was used for head comfort and immobilization. The scanner room was dimly lit. Participants were instructed to lie as still as possible with their eyes open for the duration of the run. Four eyes open ‘resting state’ runs were acquired in a single MR session from each participant (each 6m 5s yielding approximately 24m 14s of data after removal for T1 equilibration effects). One run was discarded for S2 due to motion yielding 18m 10s of data. A motor task was also collected but not analyzed here.

Functional imaging data were acquired using accelerated, multi-band single-shot gradientecho echo-planar imaging, with robust auto-calibration using the fast low-angle excitation echo-planar technique (FLEET-ACS; Polimeni et al. 2016) sensitive to BOLD contrast: TR 1490 ms, TE 24.0 ms, flip-angle 60°, 1.35 mm isotropic voxels, matrix 142 × 142 × 81, multi-slice 3x acceleration, GRAPPA factor 3. The slice acquisition plane was tilted towards the coronal plane until coverage of the cerebellum was achieved. A T1-weighted anatomical scan was acquired for each participant using multi-echo MPRAGE: TR 2530 ms, TE 1.76, 3.7 ms, TI1100ms, flip angle 7°, 0.75 mm isotropic voxels, matrix 320 × 320 × 224, GRAPPA factor 2.

### 3T Data Processing

A custom in-house pre-processing pipeline (‘iProc’) optimized within-subject data alignment across different scanning sessions, preserving anatomical detail as much as possible by minimizing spatial blurring and multiple interpolations (expanding on Braga and Buckner 2017; Yeo et al. 2011; Poldrack et al. 2015). Each subject’s data were processed separately. To optimize alignment, two subject-specific registration templates were created: a mean BOLD template and a T1 native-space template (as described below). Spatial alignment was achieved by calculating five transformation matrices to be applied to each usable BOLD volume. For each BOLD volume, three transforms were calculated to 1) correct for head motion, 2) correct for geometric distortions caused by susceptibility gradients (using the acquired B0 fieldmap), and 3) register the BOLD volume to the within-subject mean BOLD template. Two further transforms were calculated once for each subject and applied to all registered volumes. These transforms projected data 4) from the mean BOLD template to the T1 native-space template, and 5) from the T1 native-space template to the MNI ICBM 152 1mm atlas (Mazziotta et al. 1995). The transformation matrices were composed into a single matrix that was applied to the original BOLD volumes in a single interpolation to reduce spatial blur. BOLD data were projected to native space by composing matrices 1–4 and to MNI space by composing matrices 1–5. Registration details are described below. Data were also projected to the cortical surface and prepared for functional connectivity as detailed below.

#### Within-subject Spatial Alignment

Motion correction was achieved by aligning all volumes in a run to the middle volume of that run (matrix 1; linear registration using MCFLIRT; FSL v5.0.4; Jenkinson et al. 2002). Distortion correction of the middle volume was achieved using the fieldmap collected in that same MR session, yielding a non-linear transformation matrix (matrix 2; calculated using FSL FUGUE; FSL v4.0.3; Jenkinson 2004). Visual inspection of each session’s fieldmap showed that inhomogeneities varied from session to session, hence a session-specific fieldmap was used to optimize distortion correction.

The subject-specific mean BOLD template was created iteratively. First, the distortion-corrected middle volume of the run temporally closest to the fieldmap was selected as an interim registration target. The interim target was upsampled to 1.2mm isotropic space to aid subsequent spatial alignment of BOLD volumes (using FLIRT; FSL v5.0.4; Jenkinson and Smith 2001). Next, all of the distortion-corrected middle volumes from each run were registered to this interim target using linear registration at 12 degrees of freedom (dof; using FLIRT; FSL v5.0.4; Jenkinson and Smith 2001). The aligned BOLD images were averaged, creating the within-subject mean BOLD template in 1.2mm isotropic space. In this way, BOLD data from every run contributed to the subject’s mean BOLD template, minimizing bias towards any one run or session.

Alignment between the distortion-corrected middle volume of each run and the subject-specific mean BOLD template was achieved using linear registration at 12 dof (matrix 3; FLIRT; FSL v5.0.4; Jenkinson and Smith 2001). The T1 native-space template was created by selecting a T1-weighted structural image (upsampled to 1mm isotropic space) that was visually deemed to have good pial and white matter boundary surface estimates as calculated automatically by FreeSurfer’s recon-all (Fischl et al. 1999). Projection of data from the mean BOLD template to T1 native space was achieved using linear registration at 12 dof (matrix 4; FLIRT; FSL v5.0.4). Projection of data from T1 native space to MNI space was performed using nonlinear registration (matrix 5; FNIRT; FSL v5.0.4; Andersson et al. 2010).

#### Preprocessing for Functional Connectivity

Nuisance variables (6 motion parameters plus whole-brain, ventricular and deep white matter signal) and their temporal derivatives were calculated from BOLD data in T1 native space. The signals were regressed out of both native- and MNI-space data (using 3dTproject; AFNI v2016.09.04.1341; Cox 1996;2012). This was followed by bandpass filtering at 0.01–0.1 Hz (using 3dBandpass; AFNI v2016.09.04.1341; Cox 1996; 2012). MNI-space data were smoothed using a 2mm full width at half maximum (FWHM) kernel (using fslmaths; FSL v5.0.4; Smith et al. 2004). For surface analysis, data were resampled from the native space to the fsaverage6 standardized cortical surface mesh (40,962 vertices per hemisphere; Fischl et al. 1999) and then surface-smoothed using a 2mm FWHM kernel. Data were sampled from the gray matter halfway between the white and pial surfaces using trilinear interpolation.

The iProc pipeline thus allowed for high-resolution and robustly aligned BOLD data, with minimal interpolation and signal loss, output to three final spaces: the native space, MNI space, and the fsaverage6 cortical surface. The use of the MNI and fsaverage common reference spaces allowed us to display the data using standardized orientation, coordinates and location of gross anatomical landmarks, while preserving each individual’s idiosyncratic anatomy. Data were checked during matrix calculations for registration errors and quality control. Note that, unconventionally, BOLD data were here upsampled to 1mm isotropic space (in both native and MNI spaces). This substantially increased the computational load of all preprocessing and analysis steps, but was found in preliminary analyses to produce functional connectivity maps with noticeably better-resolved regions displaying finer anatomical details that were largely confined to the gray matter. The improvement was visible even when comparing to data that was more modestly upsampled to 1.5mm isotropic space.

### 7T Data Processing

The 7T data were preprocessed using similar procedures to the 3T data with a few key differences. Notably, no fieldmaps were available so no distortion correction was applied. A subject-specific mean BOLD alignment target was created by 1) calculating the mean BOLD image for each run, prior to removal of the first 4 volumes and motion correction, 2) registering the mean images to MNI ICBM 152 1mm space (Mazziotta et al. 1995) using linear registration (FLIRT; FSL v5.0.4; Jenkinson and Smith 2001), and then 3) calculating the mean of the registered images. Functional data were then processed as follows: 1) The initial 4 volumes of each run were discarded to allow for T1-equilibration effects. 2) Head motion correction transforms (matrix 1) were calculated by registering each volume to the mean BOLD image from each run (calculated after alignment to the middle volume of each run; using MCFLIRT; FSL v5.0.4; Jenkinson et al. 2002). 3) Transforms were calculated for registration between each run’s mean BOLD image and the subject-specific mean BOLD template (matrix 2) using nonlinear registration (FNIRT; FSL v5.0.4; Andersson et al. 2010). 4) Transformation matrices 1 and 2 were composed into a single matrix that was applied to each original BOLD volume, so that motion correction and registration to the mean BOLD template could be achieved in a single interpolation step to reduce spatial blur. 5) Nuisance variables (6 motion parameters plus signals from whole brain as well as lateral ventricles and deep white matter, as hand-drawn for each subject) and their temporal derivatives were regressed out (using fsl_regfilt; FSL v5.0.4). 6) Data were band-pass filtered between 0.01–0.1 Hz and smoothed using (using 3dBandpass; AFNI v2016.09.04.1341; Cox 1996). Two subjects (S1 and S3) were smoothed at 2.5mm FWHM while the other (S2) at 3.0mm FWHM, as these kernels produced robust functional connectivity maps with minimal smooth in preliminary analyses using kernels ranging from 2– 4mm FWHM.

### Functional Connectivity Analysis

For the 3T data, functional connectivity analyses were performed using both seed-based and datadriven parcellation techniques both in the volume and on the surface. For the 7T data a seed-based technique was used in the volume.

#### 3T Surface-based Analysis

Pearson’s product moment correlations between the fMRI timeseries at each vertex were computed, yielding an 81,924 × 81,924 correlation matrix (40,962 vertices per hemisphere) for each run of BOLD data. These matrices were Fisher-transformed and averaged together yielding a within-subject across-run mean correlation matrix with high stability. This average matrix was then inverse-Fisher-transformed back to correlation values and assigned to the vertices of a cortical template created in-house by combining the left and right hemispheres of the fsaverage6 surface into the CIFTI format (as described in Braga and Buckner 2017). This allowed individual vertices to be selected and the resulting correlation maps to be visualized rapidly using the Connectome Workbench’s wb_view software (Marcus et al. 2011).

For seed-based analysis in each subject, individual candidate seed vertices were manually selected from the left dorsolateral pre-frontal cortex (DLPFC) to target Networks A and B (as described in Braga and Buckner 2017). Vertices were selected to produce maps that had high correlation values and spatial distributions representative of the two networks. Specifically, the precise locations of regions were examined in different cortical zones, along the posterior and anterior midline, the lateral temporal cortex, and the medial temporal lobe (MTL), where the two networks were easily differentiated (see description of network features in the Results section). For final visualization of seed-based connectivity maps, correlation values were converted to z(r) using the Fisher-transform.

For automated parcellation analysis in each subject, *k*-means clustering was implemented. Preprocessed data were concatenated in time and MATLAB’s *kmeans* function (v2015b; MathWorks, Natick, MA) was used to parcellate the timeseries into 12 clusters on the surface (and also in MNI space). Due to the computational load, default settings (1 random initialization, 1 replication) were used that, although likely to contain local minima, provided proof of principle that data-driven parcellation could be used to identify the two networks. As the results will reveal, highly similar network estimates were found for both the seed-based and parcellation approaches suggesting that the discoveries are robust to the exact network discovery method applied.

#### 3T Volume-based Analysis

MNI-space data were analyzed using AFNI’s InstaCorr (Cox 1996; 2012; Cox and Saad 2010). 3dSetUpGroupInCorr calculated the Pearson’s product moment correlation coefficients between all pairs of voxels within a brain mask for each BOLD run. The brain mask was used to exclude non-brain voxels to reduce the computational load. The matrices were Fisher-transformed before averaging across runs using 3dGroupInCorr. This platform allows interactive selection of individual voxels and rapid visualization of their seed-based correlation maps.

The surface-defined seed vertices selected to target Networks A and B (above) were used to identify the seed voxels in the volume. To obtain MNI coordinates corresponding to the surface seed vertices, the position of each x-, y- and z-slice in MNI space was encoded with an ascending integer, creating 3 reference volumes. These reference volumes were projected to the fsaverage6 surface by first applying the inverse of each subject’s native-space-to-MNI-space transformation matrix, then projecting the reference volumes to the surface using the same sampling procedure used in data preprocessing (mri_vol2surf; Free-Surfer; Fischl et al. 1999). The exception was that both these transformations were performed using nearest neighbor interpolation to maintain the xy- and z-axis coordinates as integers. This produced three surface maps where each vertex contained a value corresponding to that vertex’s location along x-, y- or z-axes in MNI space. In this way, the MNI coordinates corresponding to the selected surface vertices for Networks A and B were used to define the volume maps.

#### 7T Volume-based Analysis

Native-space data were analyzed using AFNI’s InstaCorr as described above. The within-subject mean BOLD template was used as an underlay for anatomical guidance. At this resolution (data acquired at 1.35mm and upsampled to 1mm isotropic space), the gyral and sulcal anatomy could be observed in the mean BOLD template, providing anatomical detail with the same distortion profile as the actual BOLD data used to calculate functional connectivity. Anatomical landmarks were identified by comparison with a reference atlas containing brain images and anatomical labels at different slices and orientations (Duvernoy 1999). Individual seed voxels were selected from the gray matter along the DLPFC of each subject at or near the superior frontal sulcus and superior frontal gyrus, and the resulting correlation maps were visualized at a threshold of z(r) > 0.2. The seed voxels yielding the best estimates of the two networks were determined by checking the anatomical locations of regions in different cortical zones, along the posterior and anterior midline, the MTL (for Network A), and the inferior frontal gyrus (for Network B), where the two networks were easily differentiated (see description of network features in the Results section).

#### Overlap Maps

To visualize the spatial relationships between the two networks, overlap maps were created by thresholding the correlation maps at z(r) > 0.2 and binarizing. The binarized Network B map was multiplied by a scalar value of 2, and the binarized Network A map was subtracted from the scaled Network B map. This resulted in an image where Network A-specific voxels had a value of −1, Network B-specific voxels had a value of +2, and the overlap voxels had a value of +1. The overlap maps were visualized in wb_view (Marcus et al. 2011) using the FSL colorbar set to range from −5.0 to 0 and +0.5 to +2.5.

### Experimental Design and Statistical Analysis

This study includes n = 5 participants, two of which were scanned over 31 fMRI sessions at 3T and three of which were scanned in one session at 7T. Data were averaged over four independent datasets each including n = 12 fMRI runs to test the reliability of the connectivity patterns in the 3T study. The large volume of data allowed three independent replications within each subject. Data were averaged over n = 59 and n = 53 fMRI runs for datadriven clustering and averaged over the four replication datasets (n = 48) for seed-based best estimates of connectivity patterns in the 3T study. Data were averaged over n = 3 and n= 4 fMRI runs in the 7T study (details in ‘Functional Connectivity Analysis’ section above). Functional connectivity between brain regions was calculated in MATLAB (version 2015b; http://www.mathworks.com; Math-Works, Natick, MA) using Pearson’s product moment correlations and Fisher’s r-to-z transformation prior to averaging across runs. Network parcellation was performed using MATLAB’s *kmeans* function (version R2015b).

## Results

### Parallel Distributed Networks Are Identified Within Individuals

Networks A and B within the canonical group-averaged DN could be identified in all five individuals (Figs. 1–4). In each individual, the networks recapitulated previously described features, in particular the distributed nature of both networks, the closely juxtaposed regions observable in multiple zones of the cortex, and the close interdigitated arrangement of the two networks (Braga and Buckner 2017). Specifically, Networks A and B are distributed within canonical DN regions in the posterior medial cortex including posterior cingulate (PCC) and retrosplenial cortex (RSC), the anterior medial cortex extending throughout medial prefrontal cortex (MPFC) including anterior cingulate cortex (ACC), the inferior parietal lobule (IPL) extending rostrally to the temporoparietal junction (TPJ), and the lateral temporal cortex (extending the length of the superior temporal sulcus and middle temporal gyrus).

There were multiple anatomical details that differed between the networks. The two most prominent differences, perhaps to be considered diagnostic features that can be used heuristically to distinguish the networks, are: (1) Network A shows strong connectivity with a region of posterior PHC while Network B does not, and (2) Network B shows connectivity with rostral regions of the IPL extending into TPJ, while Network A generally shows connectivity with more caudal regions of the IPL.

**Figure 1:**
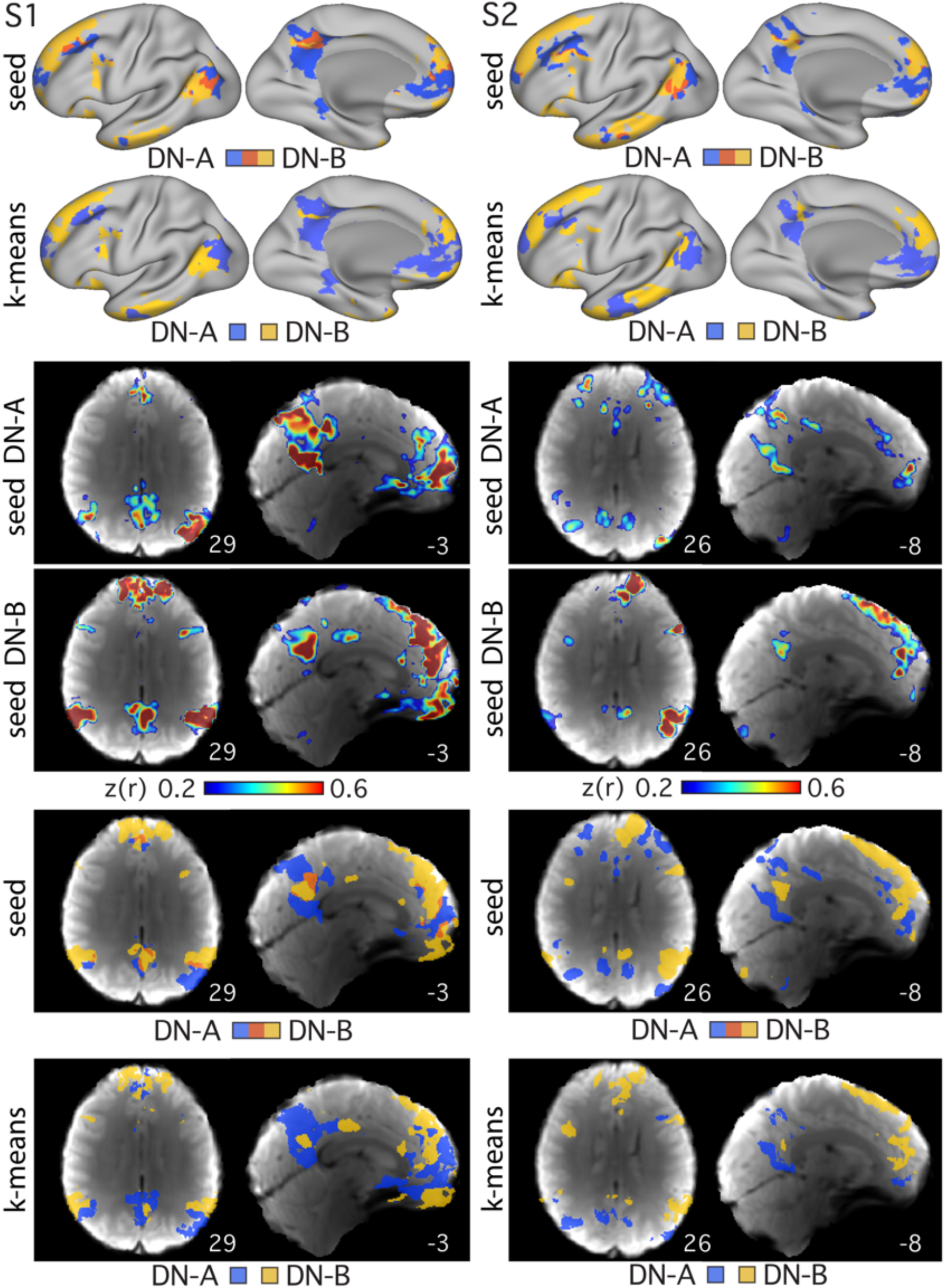
Networks are reproduced using multiple analysis methods. Data are shown on the surface (top two rows) and volume (remaining rows). Each column shows maps produced using all data from S1 (left columns) and S2 (right columns). The two networks, Network A (DN-A; in blue) and Network B (DN-B; in yellow), were defined using seed-based correlation and data-driven parcellation (*k*-means clustering at k=12). The ‘seed DN-A’ and ‘seed DN-B’ rows show the individual seed-based correlation maps from seeds in the DLPFC. The overlap images (‘seed’ rows) show the same seed-based connectivity maps thresholded at z(r) > 0.4 for S1 and z(r) > 0.25 for S2, and then binarized, with regions of overlap displayed (in red). Numbers cor-respond to MNI coordinates for each slice. Left hemisphere is on the left of each axial slice.

The relative positions of the two networks within the many canonical DN regions revealed further reliable differences, including:(3) Network A occupies more ventral regions of the posterior midline, including those at or adjacent to the RSC and ventral PCC, while Network B is typically located in a more focal dorsal region of the PCC sparing the RSC, (4) Network B involves a small circumscribed region of the ventromedial prefrontal cortex that sits beneath the regions showing connectivity with Network A, (5) Network B occupies a large swath of lateral temporal cortex that extends along the majority of the middle temporal gyrus and includes the temporal pole, while Network A occupies smaller circumscribed regions in rostral lateral temporal cortex, and (6) Network B contains a large region covering multiple gyri along the lateral inferior frontal cortex, while Network A contains only a small region more dorsally near the inferior frontal sulcus. These details replicated across the four independent datasets in both 3T study subjects (not shown), and in the three 7T study subjects (Figs. 2–4), building confidence in the separation between Networks A and B. A further analysis confirmed that the effect of TMS stimulation on the topography of functional connectivity patterns was negligible (not shown).

### Networks Are Observed Using Multiple Analysis Approaches

A potential concern was that the manual seed-selection process may have introduced observer bias towards correlation patterns that confirm Network A and Network B. To address this concern, we performed data-driven *k*-means clustering to parcellate the timeseries from all surface vertices and gray matter voxels into discrete networks. The “k-means” panels in Figure 1 show that data-driven clustering also delineated both Networks A and B in both subjects (S1 and S2). This confirms that the observation of the two networks is not dependent on the specific method of analysis.

Another possible concern surrounds distortions that might arise from surface projection. Specifically, the surface sampling procedure can induce breaks in the spatial pattern of activations by splitting a single contiguous region in the volume into multiple regions on the surface. An interdigitation pattern might thus arise artificially as a consequence. Figure 1 shows the two networks, Network A (in blue), Network B (in yellow), and their overlap (in red), defined on the surface and in the volume. The two networks occupy closely juxtaposed regions throughout the brain in the volume and show notable interdigitation of regions particularly along the posterior and anterior midline, indicating that the interdigitation was not contingent on the surface projection step. Similar features were also observed in the volume in the higher resolution data (Figs. 2–4). The replication and generalized observation of distinct Networks A and B through multiple forms of data acquisition, preprocessing and analysis techniques provide strong evidence that they represent real features of functional organization.

**Figure 2:**
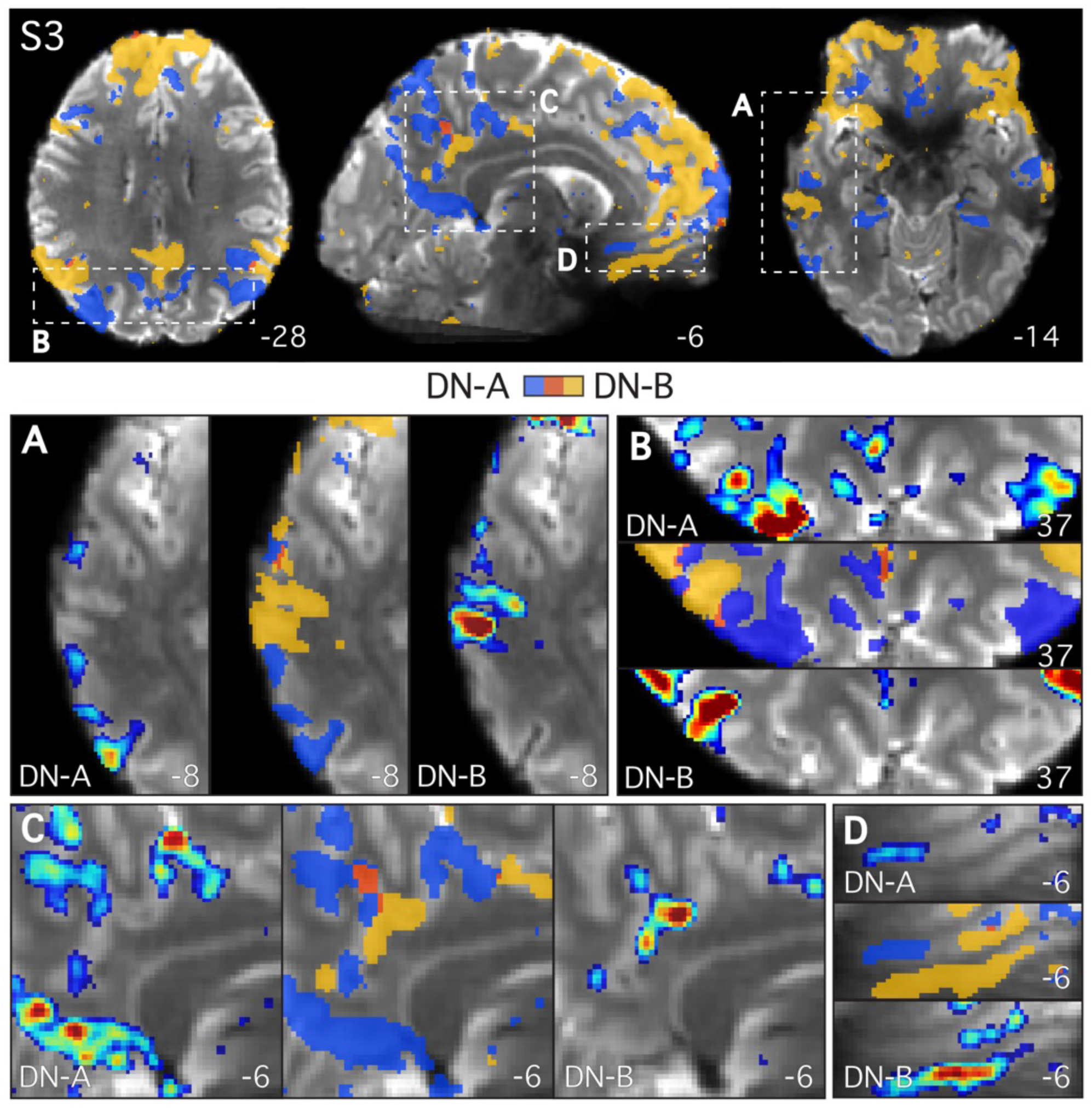
Parallel closely juxtaposed distributed networks revealed at high resolution in S3. Both Network A (DN-A; in blue) and Network B (DN-B; in yellow) were defined in S3 (7T study) at high-resolution using seed-based functional connectivity. Individual seed voxels were selected from the left DLPFC (seeds not shown). Functional connectivity maps are thresholded at z(r) > 0.2 in all images, and binarized in the ‘overlap images’ to show both networks and their overlap (in red). Note that even at this liberal threshold minimal overlap is observed. The sub-ject’s mean BOLD image is displayed as the underlay to allow the gray matter (lighter gray) and white matter (darker gray) anatomy and corticospinal fluid (white) to be seen. The regions closely follow the curvature of the gray matter and are interdigitated in complex ways along the cortical ribbon. The top panel contains full axial and sagittal slices to show the distributed par-allel organization of the two networks. The remaining panels zoom in on the lateral temporal cortex (A), the parietal lobes (B), and the posterior (C) and ventral anterior midline (D) to high-light the close juxtaposition of the regions. Functional connectivity maps are displayed using the colorbar range (0.2 – 0.6) as shown in Fig. 1. Numbers correspond to MNI coordinates for each slice. Note that different slices are sometimes shown in the insets and top panel. Left hem-isphere is on the left of each axial and coronal slice in all figures.

**Figure 3:**
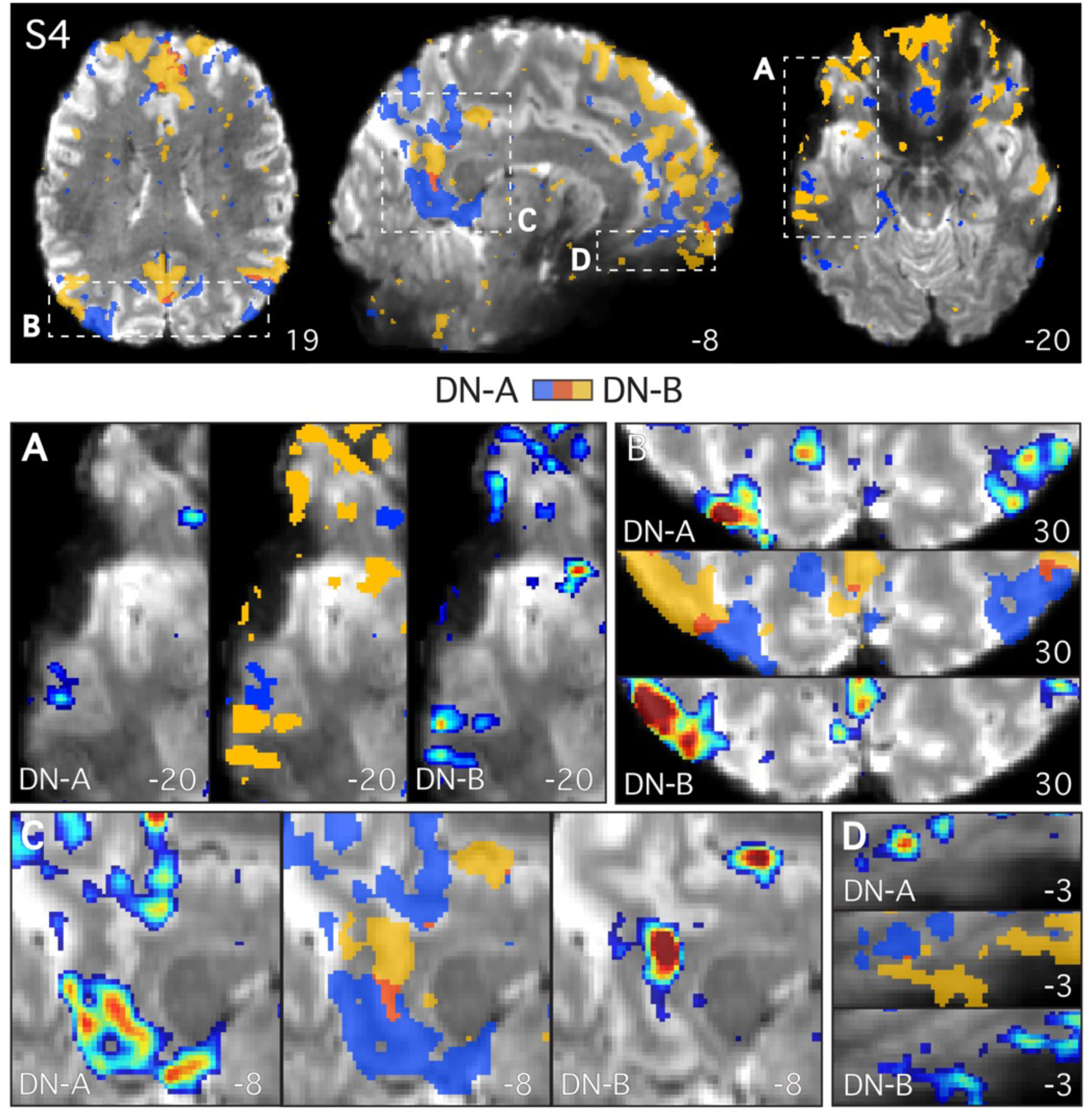
Parallel closely juxtaposed distributed networks revealed at high resolution in S4. Analysis of S4 (7T study) recapitulated features observed in S3 (Fig. 2). The two networks, Network A (DN-A) and Network B (DN-B), tightly follow the gray matter anatomy and are closely juxtaposed in multiple cortical zones. Figure formatted according to Fig. 2.

**Figure 4:**
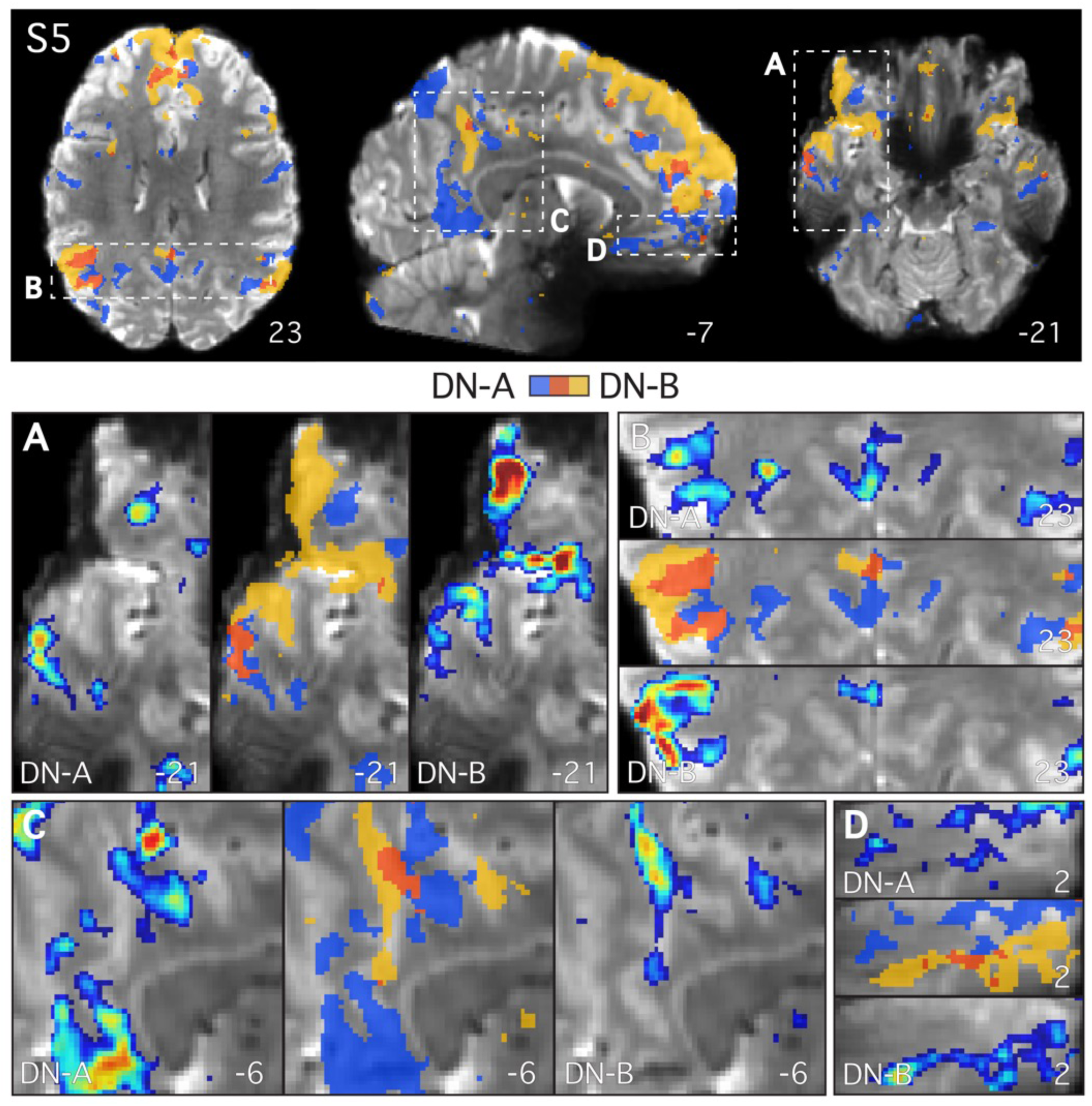
Parallel closely juxtaposed distributed networks revealed at high resolution in S5. Analysis of S5 (7T study) recapitulated detailed features observed in S3 (Fig. 2) and S4 (Fig. 3), The two networks, Network A (DN-A) and Network B (DN-B), again follow the gray matter anatomy and are closely juxtaposed in multiple cortical zones. This individual displays more overlap than the others at this threshold. Figure formatted according to Fig. 2.

### Closely Juxtaposed Networks Follow the Cortical Ribbon at High Resolution

The 7T data allowed the detailed anatomy of the distributed networks to be resolved at high resolution. Figures 2–5 zoom in on different zones of the DN so that the locations of the regions along the cortical ribbon can be appreciated. The two networks closely followed the complex geometry of the gray matter (shown as light gray pixels in Figs. 2–5). Note that in these images the white matter (dark gray pixels in Figs. 2–5) has not been masked out. A number of features were revealed when considering the networks at this spatial scale.

First, the two networks often contained regions that were closely juxtaposed along the cortical mantle, sometimes with sharp transitions between the networks (e.g., panel C in Figs. 2 and 3). Second, the two networks showed few regions of overlap in all three 7T subjects, despite the low threshold of z(r) >0.2 which permits weak correlations (< 0.3) to be observed. Third, the two networks can occupy distinct regions deep within the same sulcus, sometimes on opposite banks a few millimeters apart (panels A, C, F, G and H in Fig. 5). Fourth, regions of the two networks were sometimes interdigitated in complex ways, with sequential regions following the curvature of the gray matter (panels A, C, D and F in Fig. 5). Fifth, a region belonging to one network can sit deep within the fundus, and be surrounded on either side by regions belonging to the other network (e.g., panel F in Fig. 5). There did not appear to be a clear relationship between the geometry of the gray matter and the location of the transition zone between the networks. It is not the case that a specific region of one network was always situated or always extended to a specific point along the sulcus, such as the fundus (see also Amunts et al. 1999). Rather, the network regions and their transitions were observed at different points in different sulci.

The above observations illustrate the extent to which the two networks are closely intertwined within the cortical anatomy and suggest, within the resolution and sensitivity available in these data, that the networks often occupy non-overlapping regions of the cortical sheet.

**Figure 5:**
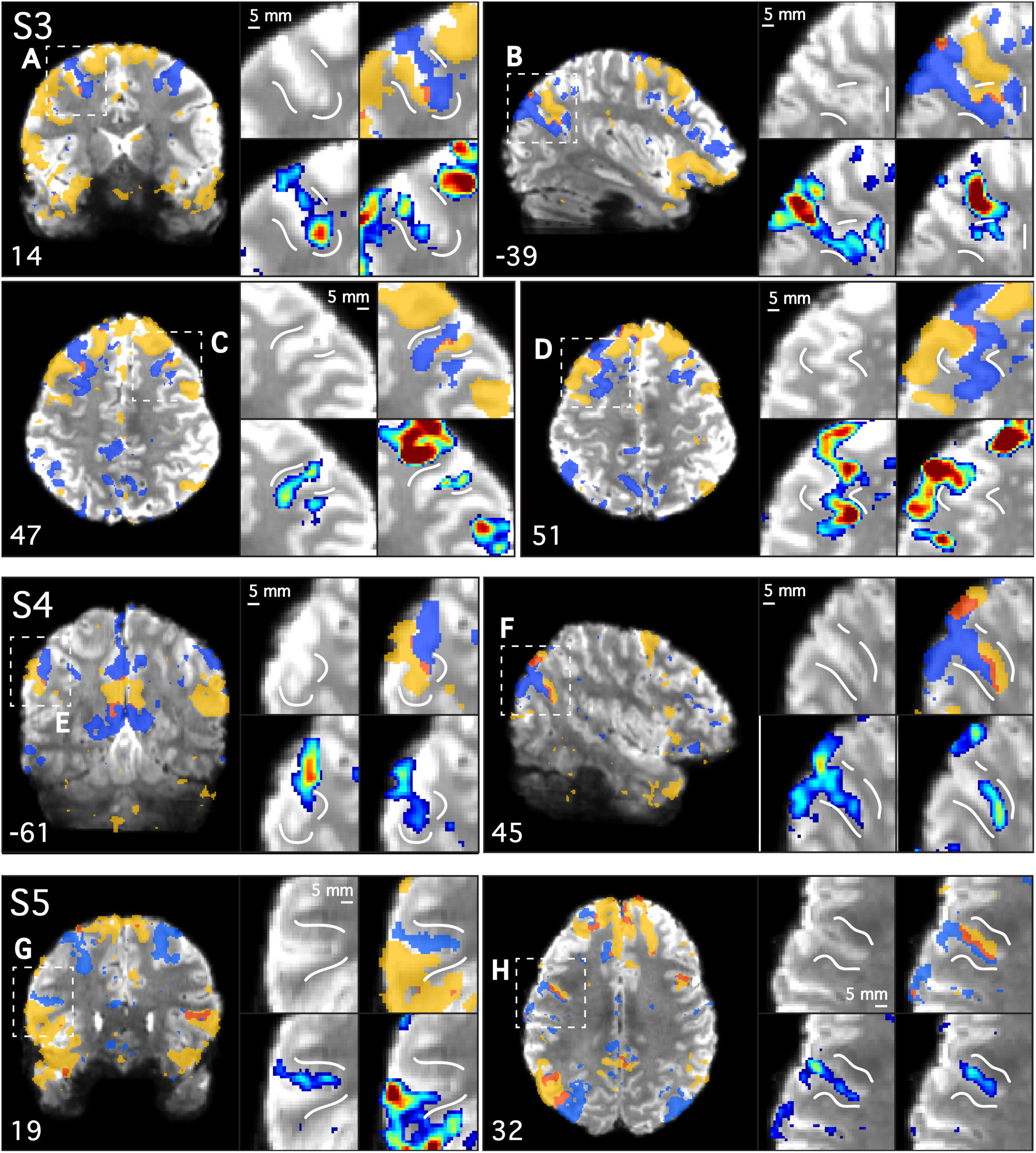
Networks are closely juxtaposed inside individual sulci. High-resolution data from three subjects are shown (S3, S4 and S5; 7T study). Selected slices illustrate locations where the two distributed networks fall in distinct regions of the same sulcus or junction between sulci. Network A (DN-A; in blue), Network B (DN-B; in yellow), and their overlap (in red) are shown. In each panel, an overlap image is shown in the full coronal (A, E, G), sagittal (B, F) or axial (C, D, H) slice, followed by four insets that zoom in on the dashed-line box to highlight anatomical details. The insets include each subject’s mean BOLD underlay for anatomical guidance (top left), the overlap image (top right), and the individual functional connectivity maps for Network A (bottom left) and Network B (bottom right). The white lines provide landmarks so that the relative position of each network can be considered across insets. The two networks were often observed on opposite banks of the same sulcus, deep into the fundus (see panels A, C, G and H). Maps are thresholded at z(r) > 0.2 in all images, and functional connectivity maps are displayed using the colorbar range (0.2 – 0.6) as shown in Fig. 1. Note other views of these same functional connectivity maps are displayed in Figs. 2–4.

### Consistent Spatial Relationships at High Resolution

Consistent spatial relationships were present when the networks were considered in all three high-resolution individuals together (Figs. 2**–**5). Such details may be informative for techniques that seek to record from or stimulate specific networks (e.g., intracranial methods). Below we describe some of these features. Sulcal and gyral anatomy were determined by comparison with the Duvernoy (1999) reference atlas. The observations are intended to provide approximate anatomical landmarks.

#### Posteromedial Cortex

In the posterior midline (panel C in Figs. 2–4), the two networks occupied sequential regions in an interdigitated fashion, beginning in the cingulate sulcus at the most anterior portion of the PCC, and descending through the subparietal sulcus into ventral PCC and RSC. The callosal sulcus (thin light-gray band circling the corpus callosum in the top panel of Figs. 2–4) was spared by both networks even in the region immediately posterior to the splenium (note that in S3 in Fig. 2 Network A skirts the RSC border immediately inferior to where the callosal sulcus terminates). The interdigitated network regions extended into the marginal segment of the cingulate sulcus, containing a Network A region at the junction between the subparietal and cingulate sulci in all three subjects. More ventrally, the interdigitated regions also extended into the transverse parietal sulcus (which originates near to the subparietal sulcus and ascends to the dorsal precuneus, bisecting the medial parietal lobe), containing a large Network B region at or immediately ventral to the junction with the subparietal sulci in all three subjects.

A consistent feature was that Network A occupied the most ventral regions of posteromedial cortex, while Network B generally occupied dorsal regions of the PCC. Specifically, a large region of Network A was found to extend along the dorsal bank of the parieto-occipital fissure in all three participants. Network A also consistently occupied more dorsal regions in the posterior midline (i.e. the dorsal precuneus). In this way, Network A could be seen to surround on three sides the Network B region in the PCC.

Finally, following the cingulate sulcus rostrally, side-by-side regions could be seen extending as far as the middle cingulate cortex (Vogt et al. 2009) in all three subjects, confirming the previous observation of this zone as a site of juxtaposition between the two networks (see Fig. 3 ‘zone 6’ in Braga and Buckner 2017).

#### Medial Temporal Lobe

An anatomical difference previously observed between the two distributed networks was that Network A exhibited strong correlation with a region of the MTL at or near the posterior PHC, whereas no such evidence was found for Network B (even in more a focused analysis; see Fig. S2 in Braga and Buckner 2017). In the present high-resolution data we explored the functional connectivity of the MTL in the volume in more detail, to ascertain if further insights could be gained about the relation of this region to Networks A and B.

Network A exhibited strong correlation with a region of posterior PHC in all three subjects. This region lay deep within the collateral sulcus (see top left axial views in Figs. 6 and 8) and extended along the posterior-anterior axis of this sulcus, beginning just anterior to the atrium of the lateral ventricles and extending rostrally. It was difficult to ascertain the most anterior extent of this PHC region due to this section of the MTL being affected by magnetic susceptibility artifacts from the nearby ear canals and sinus (Ojemann et al. 1997). Thus, it is unresolved whether the Network A region extends further or, alternatively, whether there is representation of Network B in this zone which includes entorhinal cortex.

In all three participants, an additional region of Network A could be resolved at or near the crown of the subiculum within the hippocampal formation itself further demonstrating this network’s inclusion of traditional declarative memory structures. Panels A and B in Figures 6–8 zoom in on the MTL in each subject on a coronal slice to highlight the anatomical details of this subicular region. The region can be observed near or at the crown of the subiculum, distinct from the PHC region in the collateral sulcus and further into the hippocampal formation. To confirm the validity of this region as a component of Network A, a seed voxel was placed directly in the subiculum, and the resulting correlation map recapitulated the full and selective distribution of Network A (lower panel in Figs. 6–8). The subicular region was situated towards the middle of the anterior-posterior hippocampal axis, approximately where the MTL curves around the brainstem, at the level where the raphe nuclei can be observed in the axial plane. Depending on the individual’s anatomy, the subicular and PHC region could be observed on the same axial and coronal slices (see Figs. 6–8).

The functional connectivity of the hippocampus was also explored at lower thresholds (z(r)≈0.15). Hints of small regions belonging to Networks A and B could be observed at these thresholds, particularly in the subject that provided the most robust data (S1). However, in the remaining subjects, these hippocampal regions were suggestive only and could not be conclusively established. Further analyses of even higher-resolution and higher-signal-to-noise data may be able to resolve regions within the hippocampal formation that show connectivity with the distributed association networks.

**Figure 6:**
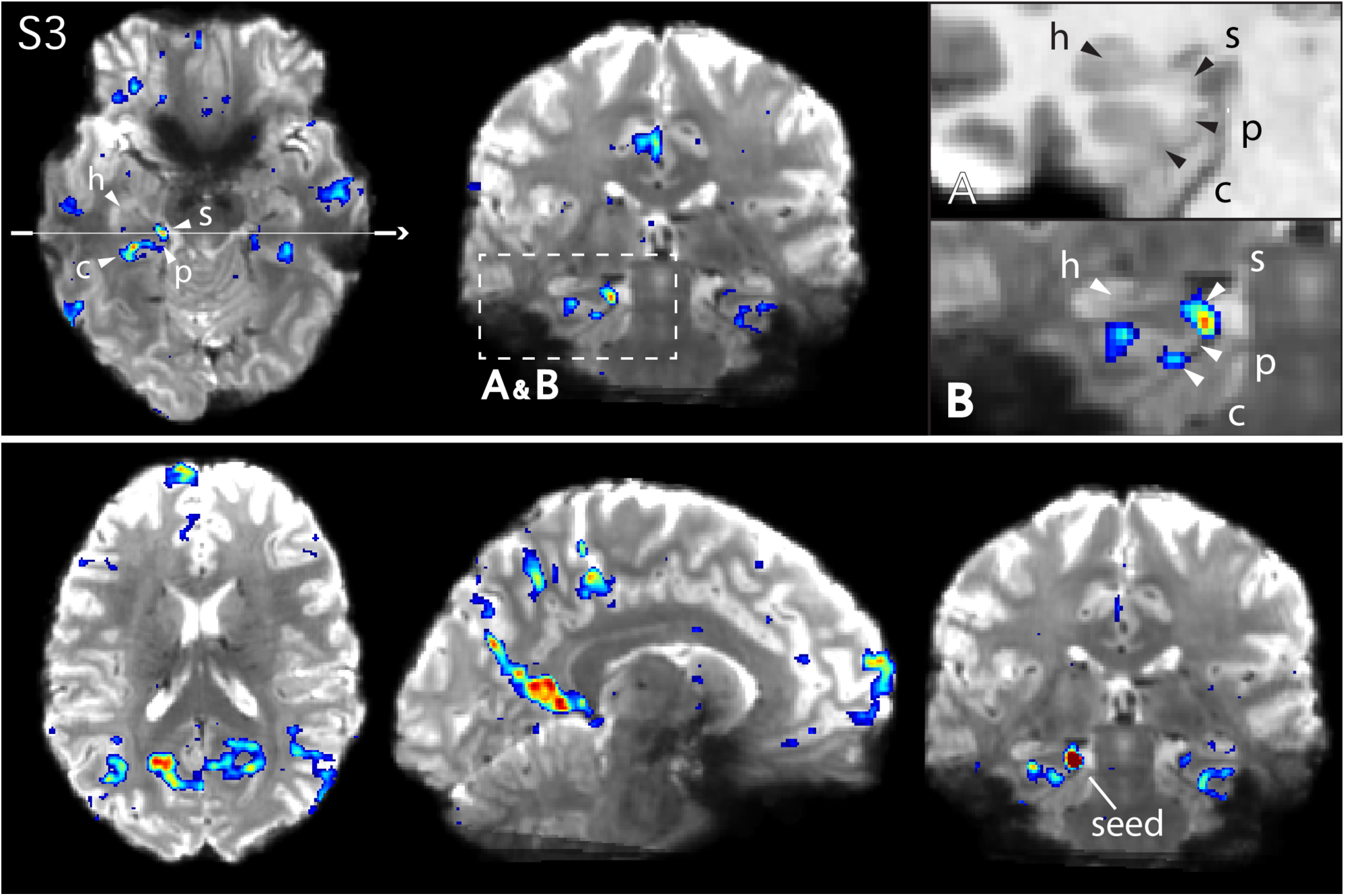
Network A shows connectivity with circumscribed regions of the medial temporal lobe in S3. The top panel shows Network A in S3 defined using a left DLPFC seed (not shown). Arrows on the axial slice (left) and insets denote the approximate locations of the hippocampus proper (h), the subiculum (s), the parahippocampal cortex (p), and the contralateral sulcus (c). The thin white line denotes the position of the coronal slice (middle). Insets show magnified images of the MTL in the coronal slice for the subject’s anatomical (T1-weighted) image (A), to display the white and gray matter anatomy clearly, and the functional connectivity map for Network A overlaid onto the subject’s mean BOLD image (B). The mean BOLD underlay allows regions to be visualized in relation to the true BOLD anatomy given image distortion. Note that the arrows are placed at slightly different locations in insets A & B to match the anatomy in each image. The lower panel displays the functional connectivity map from a seed placed directly in the subiculum, which recapitulates the cortical components of Network A including the frontal pole near BA10 (see Fig. 2 for comparison).

**Figure 7:**
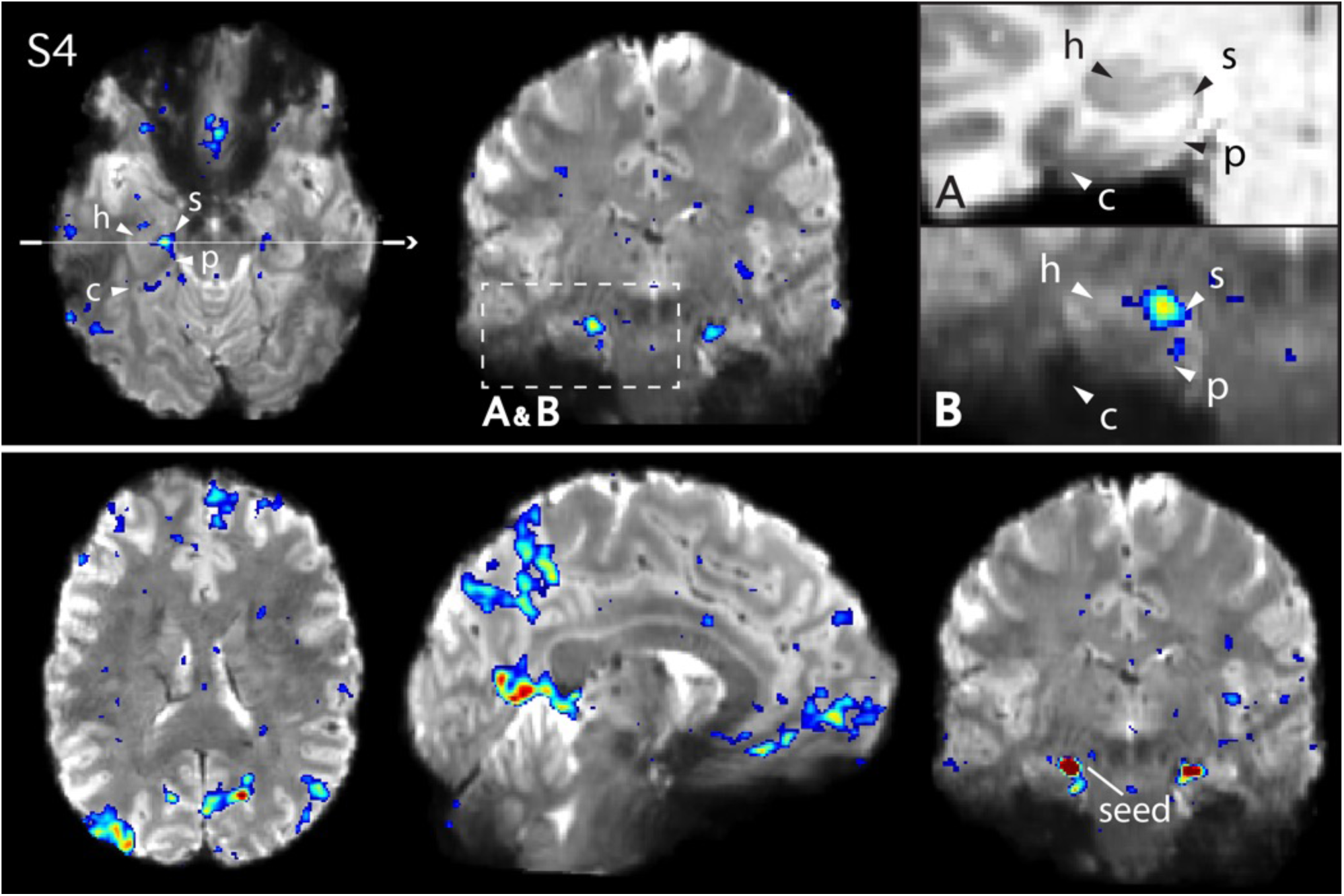
Network A shows functional connectivity with circumscribed regions of the medial temporal lobe in S4. Figure formatted according to Fig. 6 but showing data from S4. Arrows denote the approximate locations of the hippocampus proper (h), the subiculum (s), the parahippocampal cortex (p), and the contralateral sulcus (c). See Fig. 3 for comparison with similar views of the network defined from the DLPFC seed.

**Figure 8:**
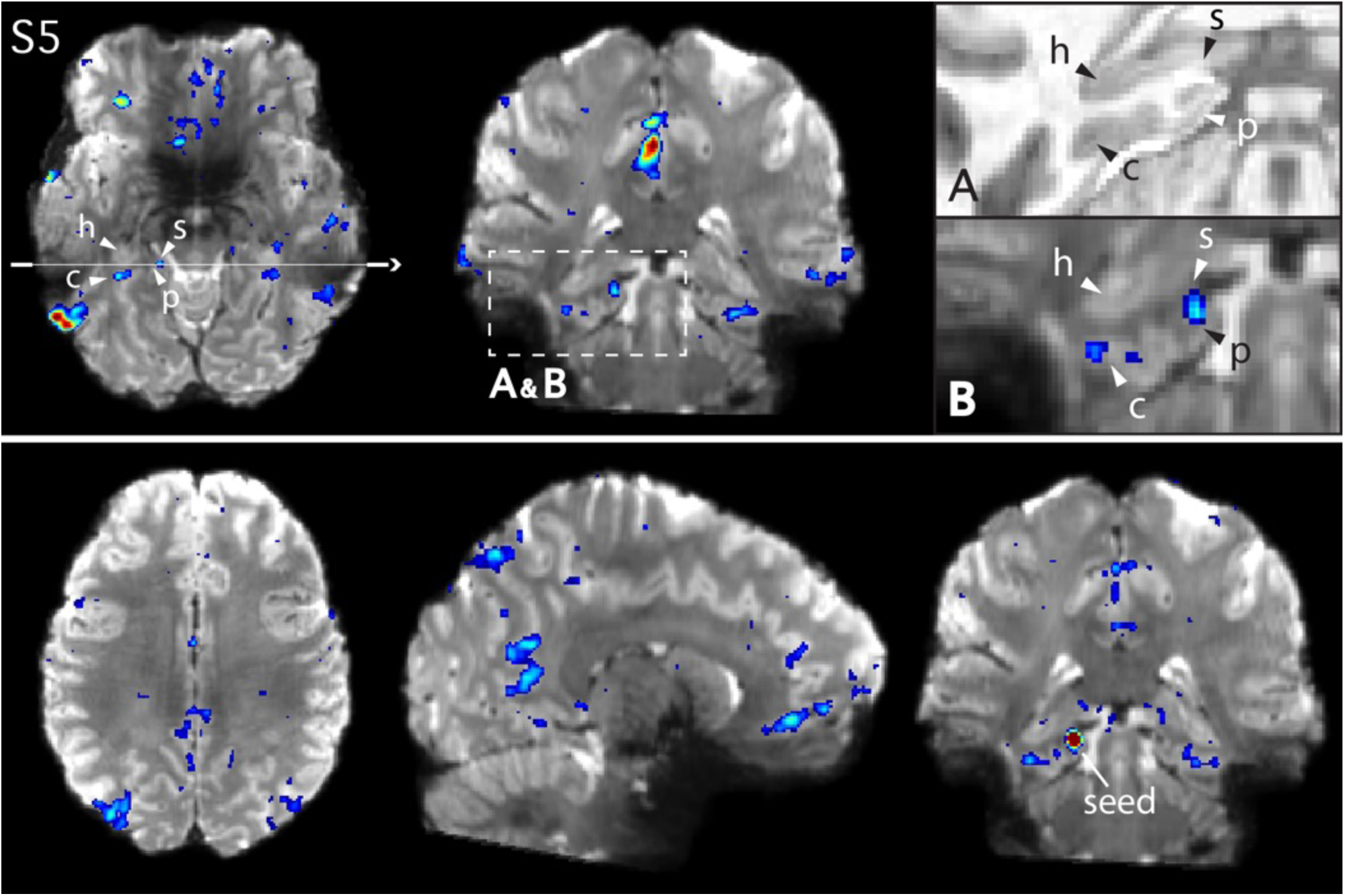
Network A shows functional connectivity with circumscribed regions of the medial temporal lobe in S5. Figure formatted according to Fig. 6 but showing data from S5. Arrows denote the approximate locations of the hippocampus proper (h), the subiculum (s), the parahippocampal cortex (p), and the contralateral sulcus (c). See Fig. 4 for comparison with similar views of the network defined from the DLPFC seed.

#### Medial Prefrontal Cortex

The two distributed networks were closely interdigitated and occupied alternating regions in the MPFC (sagittal view in top panel in Figs. 2–4 and panel D in Fig. 2). In general, Network B occupied a large swath of MPFC in all subjects. Network B regions extended from the dorsal parts of the MPFC, descending rostrally along the midline, and reaching to the most inferior portions of the MPFC. Network A regions were interdigitated with Network B regions across the MPFC in a complex organization, though generally did not extend as dorsally or ventrally as Network B regions. A Network A region was identified in the most rostral portion of the MPFC, at or near the medial aspect of area 10 as originally demarcated by Brodmann (Fig. 86 in Brodmann 1909/2006; see updated architectonic descriptions in Petrides and Pandya 1994; Öngür et al. 2003).

The ventral portion of the anterior midline is a site of projections from limbic structures, including the amygdala (Öngür and Price 2000) and hippocampal formation (Rosene and Van Hoesen 1977). An important observation is that a Network B region was located at a more ventral region of the anterior midline than Network A. This was surprising because, in contrast with Network A, Network B does not show a strong correlation with posterior PHC, although coupling to anterior MTL regions where artifact and signal loss are prominent cannot be ruled out. Network A occupied a dorsal subgenual region, while Network B occupied more inferior regions extending rostrally along ventral MPFC (sagittal view in top panel in Fig. 2 and panel D in Figs. 2–4; see Fig. 5 in Córcoles-Parada et al. 2017, for sulcal variations in this region). Although this region is susceptible to signal drop out in fMRI, a subgenual region belonging to Network A was resolved in all three individuals (sagittal view in top panel of Figs. 2–4).

## Discussion

The canonical DN is comprised of at least two parallel distributed networks when defined within the individual (Braga and Buckner 2017). The two networks, referred to as Networks A and B, display closely neighboring regions in multiple zones of the cortex and follow a general motif characteristic of the association networks (Goldman-Rakic 1988; Yeo et al. 2011; Power et al. 2011; Doucet et al. 2011; Margulies et al. 2016; Buckner and Margulies 2018). In the present study we resolved the two networks using high-resolution scanning at 7T and describe a number of novel features with particular emphasis on anterior and posterior midline regions previously hypothesized to be zones of convergence.

### Parallel Distributed Networks Within the Default Network Are Replicated

Two networks within the canonical group-defined DN were reliably observed in five new individuals (Figs. 1–4) and identified using both manually selected seed-based correlation and datadriven clustering, on the surface and in the volume (Fig. 1). These results demonstrate that the definition of the networks is not dependent on idiosyncratic analysis choices or due to experimenter bias. Further, the two networks were identified in high-resolution data at 7T (Figs. 2–4). These collective results offer strong evidence that the observation of distinct Networks A and B is reliable.

### Parallel Distributed Networks Are Closely Juxtaposed Along the Cortical Ribbon

Anatomical details of the two networks were resolved by analyzing high-resolution data. At high spatial resolution, the two networks followed the geometry of the gray matter (Figs. 2–5). The two networks were often closely juxtaposed along the cortical mantle, displaying sharp transitions between regions (see magnified images in Figs. 2–5, and especially Figs. 5E and 5G). One individual (S5; Fig. 4) generally showed more overlap than the others.

It is not possible in the present data to determine if the networks occupy truly nonoverlapping regions of the cortical sheet. Some degree of apparent overlap is expected due to the limited resolution (e.g., partial volume effects) and spatial smoothing. However, it is notable that as we increased the effective resolution, going from 3T to 7T, the emergent picture was one of more separation rather than more overlap. In some locations, minimal overlap was observed between the two networks, despite the use of a liberal threshold and no restrictions being placed on a voxel being correlated with both seeds.

The separation of the two networks along the posterior and anterior midline in the present data was striking (e.g., Figs. 2C, 3C, and 4C). Prior studies have emphasized that the posterior midline may be a ‘hub’ or ‘core’ that interacts with multiple distinct subnetworks or distributed systems (Buckner et al. 2009; Tomasi and Volkow 2010; Andrews-Hanna et al. 2010; 2014; Braga et al. 2013; see also Power et al. 2013). The present findings raise the alternative possibility that the networks contain distinct regions situated near to one another along the complex cortical geometry of the midline. In group-averaged data, these regions are blurred together giving the appearance of a network-general or task-general convergence zone.

One prior task-based study of individuals at 7T noted a complex organization of domain-specific subdivisions at or near the precuneus that responded differentially to tasks demanding orientation to space, time, and person (Peer et al. 2015). Analysis of their detailed results in light of the present observations yields an interesting point of concordance. In their study, they identified a region in the center of the precuneus responding most strongly to ‘person-oriented’ emphasis that was surrounded by three adjacent regions responding to ‘space-oriented’ emphasis (see Fig. 4A in Peer et al. 2015). This topography is quite similar to the presently observed organization that differentiates Network A from Network B. Specifically, in three of their individual subjects, a clear juxtaposition of the person-oriented response can be seen surrounded by distinct space-oriented responses (see Subjects 4, 6, and 12 in Supplementary Fig. 1 in Peer et al. 2015). A parsimonious explanation for the collective results may be that their task paradigm was differentially activating Network A for the space-oriented condition and Network B for the person-oriented condition.

In the present data, no consistent relationship was noticed between the gross cortical anatomy (i.e., the gyral and sulcal folds) and the position of transition zones between regions across individuals. In some cases, the juxtaposition between the two networks was simple, with two neighboring regions sitting side-by-side along the cortex (e.g., Figs. 5C and 5G). In other cases, the two networks displayed a complex interdigitation, with alternating regions following the cortical ribbon in an irregular pattern (e.g., Figs. 2C and 5F).

Interdigitation of closely positioned regions has been previously observed in the cortex at various spatial scales. Anatomical projections from prefrontal and parietal association zones converge on adjacent columns of the cortical mantle in the anterior and posterior midline (see Figs. 4 and 6 in Selemon and Goldman-Rakic 1988). Alternating ocular-dominance bands permeate striate cortex that display sharp boundaries between bands (Hubel and Wiesel 1979). Further along the visual hierarchy, face-responsive regions of the inferior temporal lobe appear as a set of non-contiguous islands (the “Fusiform Face Archipelago”; Kanwisher 2010; Spiridon et al. 2006; Moeller et al. 2008) surrounded by regions that respond preferentially to other visual stimulus categories (e.g., colors, scenes or objects; Lafer-Sousa and Conway 2013; Lafer-Sousa et al. 2016). While it is not presently possible to link our fMRI findings with these anatomical features, it is nonetheless intriguing that the distributed parallel arrangement of closely juxtaposed and interdigitated functional regions echoes the distributed network features observed in direct anatomical studies. One possibility is that such features emerge because they are all outcomes of competitive activity-dependent processes that sculpt cortical organization during early development.

### Distinct Regions Are Revealed Within the Medial Temporal Lobe

Network A is robustly coupled to a specific region of the MTL at or near to the posterior PHC, while Network B is not (Braga and Buckner 2017). The present analyses confirmed this distinction repeatedly (see surfaces in Fig. 1). In the highresolution data, the PHC region participating in Network A was resolved to a portion of the contralateral sulcus in all three 7T subjects (Figs. 6 and 8 for S3 and S5; other subject not shown). Comparison of Network A, estimated here using functional connectivity measures in humans, with direct anatomical data suggests that Network A may reflect anatomical projections including from the PHC (Buckner et al. 2008; Binder et al. 2009; Buckner and Margulies 2018). Of particular interest, the PHC projects to frontopolar cortex A10 in New World monkeys (Burman et al. 2011; Rosa et al. 2018). When multiple anterior and posterior tracer injections are examined in relation to one another, a distributed network emerges that has clear parallels to Network A in terms of its wide distribution as well as its specificity (Buckner and Margulies 2018). As the last common ancestor between New World monkeys and humans lived approximately 45 million years ago, these results suggest an ancient anatomical circuit motif supports the DN. An open area for future research is to understand better whether there is expansion and differentiation of this basic circuit motif (e.g., into Network A and Network B) in the large-brained primates including apes and humans that underpin our specialized cognitive capabilities.

The high-resolution data allowed us to resolve a key additional anatomical finding. A distinct region of the MTL, inside the hippocampal formation, at or near to the subiculum, also showed functional connectivity with Network A (Figs. 6–8). In one participant (S4; Fig.7) this region was positioned beyond the crest of the parahippocampal gyrus, extending to the beginning of the hippocampal sulcus. The subiculum is a source of direct cortical efferents to the RSC as well as subgenual prefrontal cortices (Rosene and van Hoesen 1977). These subicular projections may form part of the larger distributed network we delineate here using functional connectivity (see lower panels in Figs. 6–8 which show the network defined from a seed placed in the subiculum). In the present data we were also able to detect hints of smaller regions within the hippocampus of one subject, but these regions displayed low correlations with Network A and were not reliably observed across individuals. Further exploration of the MTL at higher resolution using sequences optimized to avoid signal loss in the anterior portions of the MTL will be required to resolve details and determine whether there are components of Network B within the anterior MTL.

### Technical Considerations of Fine-Scale Functional Architecture

The fractionation of the default network into parallel networks was possible because of focus on the individual rather than group-averaged data. Prior work has shown that, even after spatial normalization, the location of functional regions can differ between individuals on the order of centimeters (e.g., see Fig. 6 in Steinmetz and Seitz 1991; Fig. 12 in Rajkowska and Gold-man-Rakic 1995; Clark et al. 1996; Henssen et al. 2016). In some cases, we were able to resolve distinct regions within the same sulcus, millimeters apart (Fig. 5C). With even a modest amount of additional blurring across or within individuals it is likely that these distinctions would be lost. Thus our results illustrate the utility of focusing on individuals, not as a means to reveal individual differences, but as an approach to maintain as much spatial information as possible and allow functionally specialized circumscribed regions to be sampled accordingly.

In addition to focusing on individuals, steps were taken to minimize spatial blur in the current analyses. BOLD data were acquired using 1.35 and 2.4mm isotropic voxels, preprocessed and re-sampled to 1mm resolution using a single interpolation step, and smoothed minimally at 2–3mm FWHM. It is likely that these steps contributed to the ability to find the present distinctions. The close juxtapositions that were revealed underscore the importance of delineating brain networks at the level of the individual, with sufficient data and resolution to resolve fine-scale features of functional architecture (Fedorenko et al. 2012; Michalka et al. 2015; Laumann et al. 2015; Huth et al. 2016; Braga and Buckner 2017; Gordon et al. 2017).

There is reason to suspect that smaller network regions remain unresolved. In addition to the hints of small regions in the hippocampus discussed above, other findings were at the edge of detection in the subject that provided the best data. Although these regions included few voxels, their spatial arrangement suggested an actual biological origin. In particular, small alternating side-by-side and bilateral regions belonging to Networks A and B were observed within the callosal sulcus extending from beneath the middle cingulate gyrus to the subgenual zone (not shown). These small regions also displayed low correlations and were not reliably observed in the other two high-resolution subjects. These observations are described here only to illustrate the possibility that additional analyses at higher resolution may resolve finer scale regions, particularly in deeper cortical and subcortical zones. Put simply, we are certain to still be missing important structure, even at the level of macroscale organization, that sits beyond our present resolution, data quality, and analysis strategies.

## Conclusions

The present analyses extend our understanding of the fractionation of the DN into multiple parallel, distributed networks. The juxtaposition of the two networks adjacent to one another along the cortical ribbon and within individual sulci underscores the need for functional anatomy to be examined at the level of the individual and at sufficient resolution. Precision functional mapping could aid applied endeavors that seek to record from or stimulate specific networks (e.g., using intracranial methods) as well as those seeking to limit complications arising from surgical resection.

## Acknowledgements

R.M.B. was supported by Wellcome Trust grant 103980/Z/14/Z and NIH Pathway to Independence Award grant K99MH117226. M.C.E. was supported by Mentored Patient-Oriented Career Development Award grant K23MH099413. This work was also supported by Kent and Liz Dauten, NIH grants P50MH106435 and P41EB015896, and Shared Instrumentation Grants S10OD020039 and S10RR019371. We thank the Harvard Center for Brain Science neuroimaging core and FAS Division of Research Computing for support. L. DiNicola assisted with data processing. A. Youssoufian, H. Becker, E. Phlegar and M.K. Drews assisted in data acquisition. B.T.T. Yeo and P. Angeli assisted with the projection of MNI coordinates to the cortical surface. H. Hoke, T. O’Keefe, R. Mair and S. McMains assisted with data processing optimization. B. Keil assisted with the use of the custom 7T head coil. The multi-band EPI sequence was generously provided by the Center for Magnetic Resonance Research (CMRR) at the University of Minnesota.

## Supporting Information

**Supporting Figure S1:**
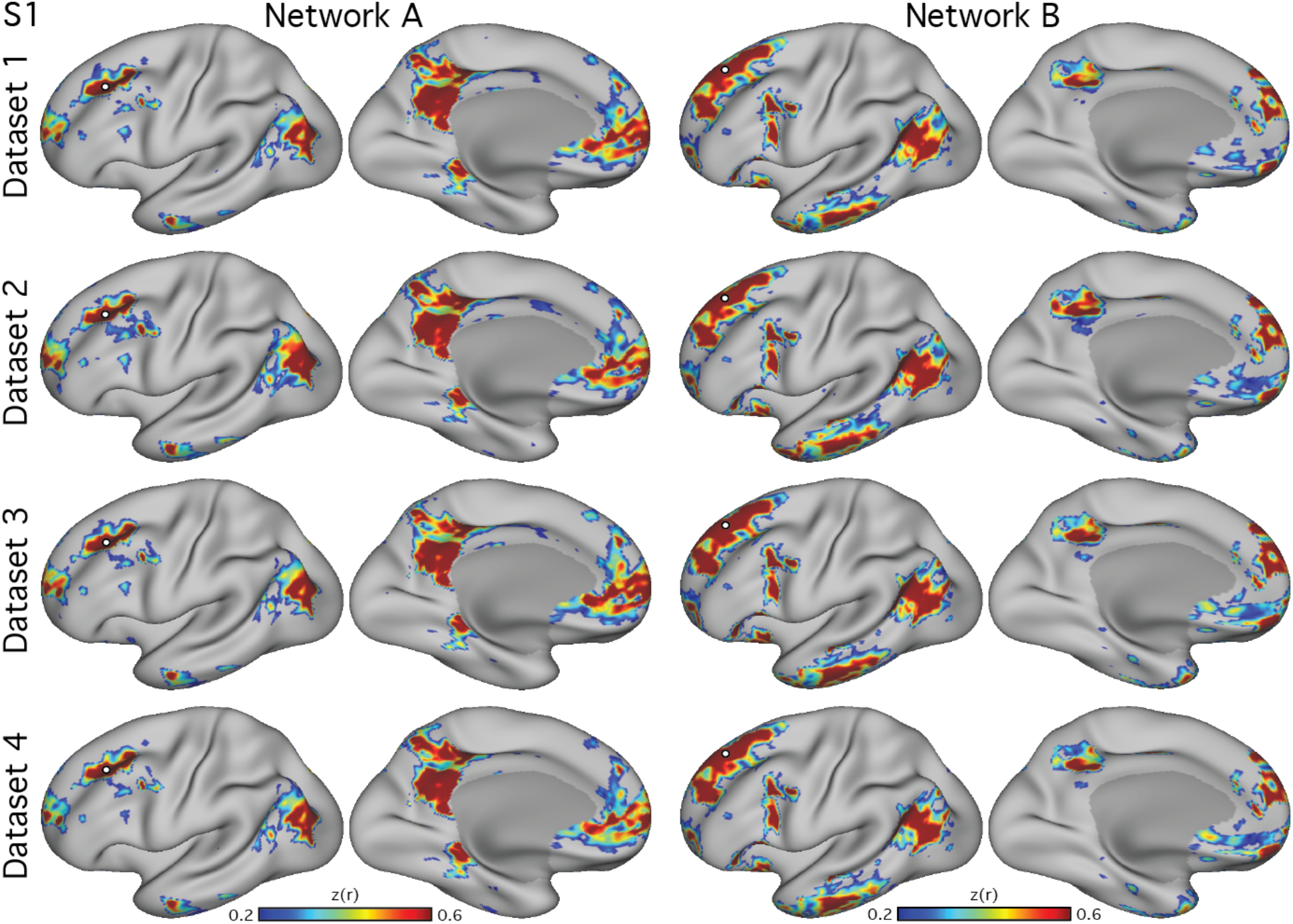
Parallel closely juxtaposed distributed networks are reliable in subject S1. Each column shows the functional connectivity maps from a single seed vertex (white circle) selected from the left dorsolateral prefrontal cortex (DLPFC) of S1 (3T study). The left pair of columns shows Network A and the right pair Network B. Data from S1 was divided into 4 independent datasets (n=12 runs or 1h 22m of data in each set). The two networks, Network A and Network B, were reliably observed in all four datasets. Each seed produced a distinct distributed network that occupied closely juxtaposed regions within zones typically considered part of the canonical ‘default network’. Lateral and medial inflated views of the left hemisphere are shown. The surfaces are rotated by 19° along the y plane to better show the ventromedial prefrontal cortex. The same view is used in subsequent figures.

**Supporting Figure S2:**
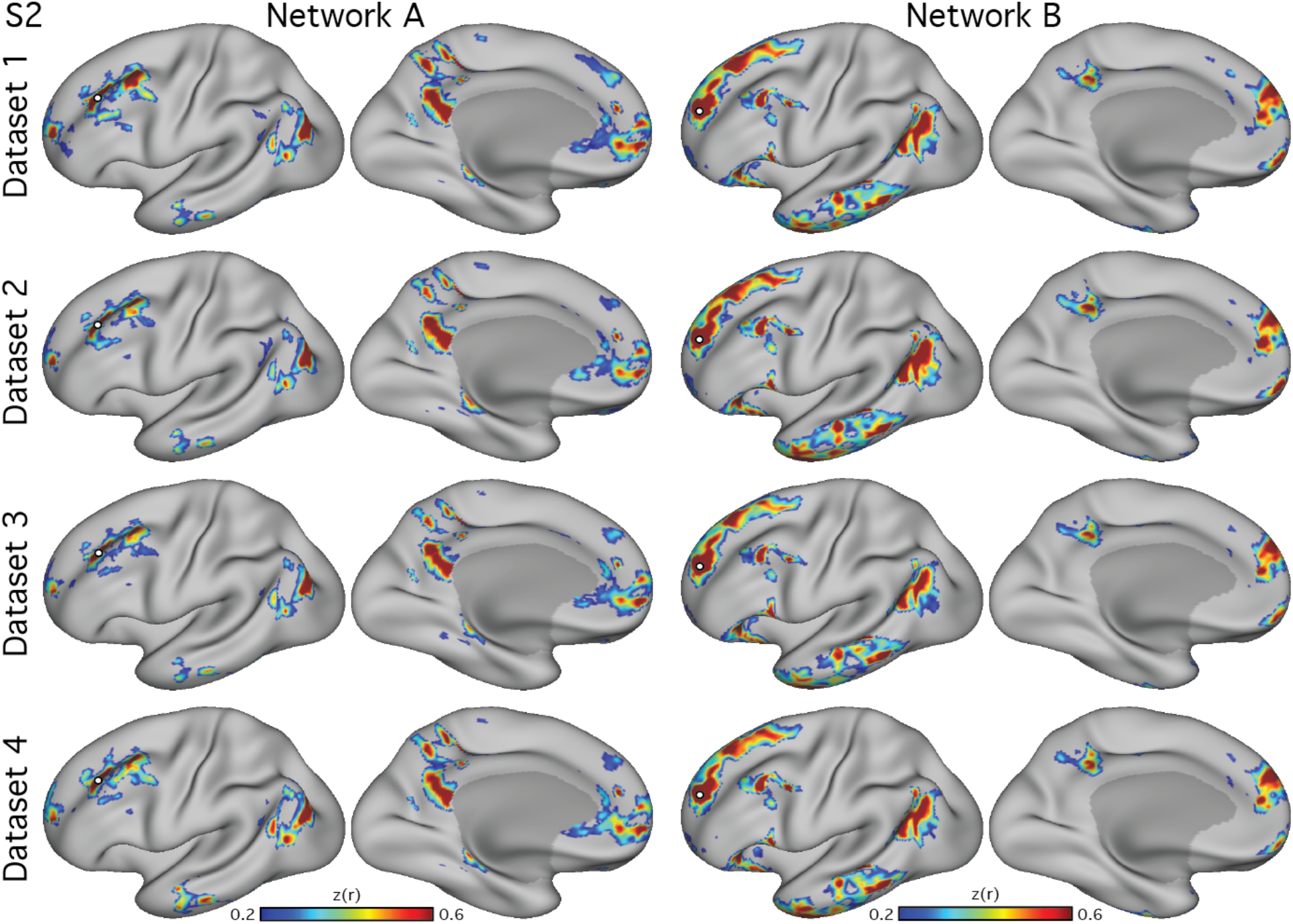
Parallel closely juxtaposed distributed networks are reliable in S2. Analysis of another individual S2 (3T study) recapitulated the observation of the two parallel interdigitated networks, Network A and Network B, consistently across four datasets (dataset 1–4). Figure formatted according to Supporting Fig. S1.

**Supporting Figure S3:**
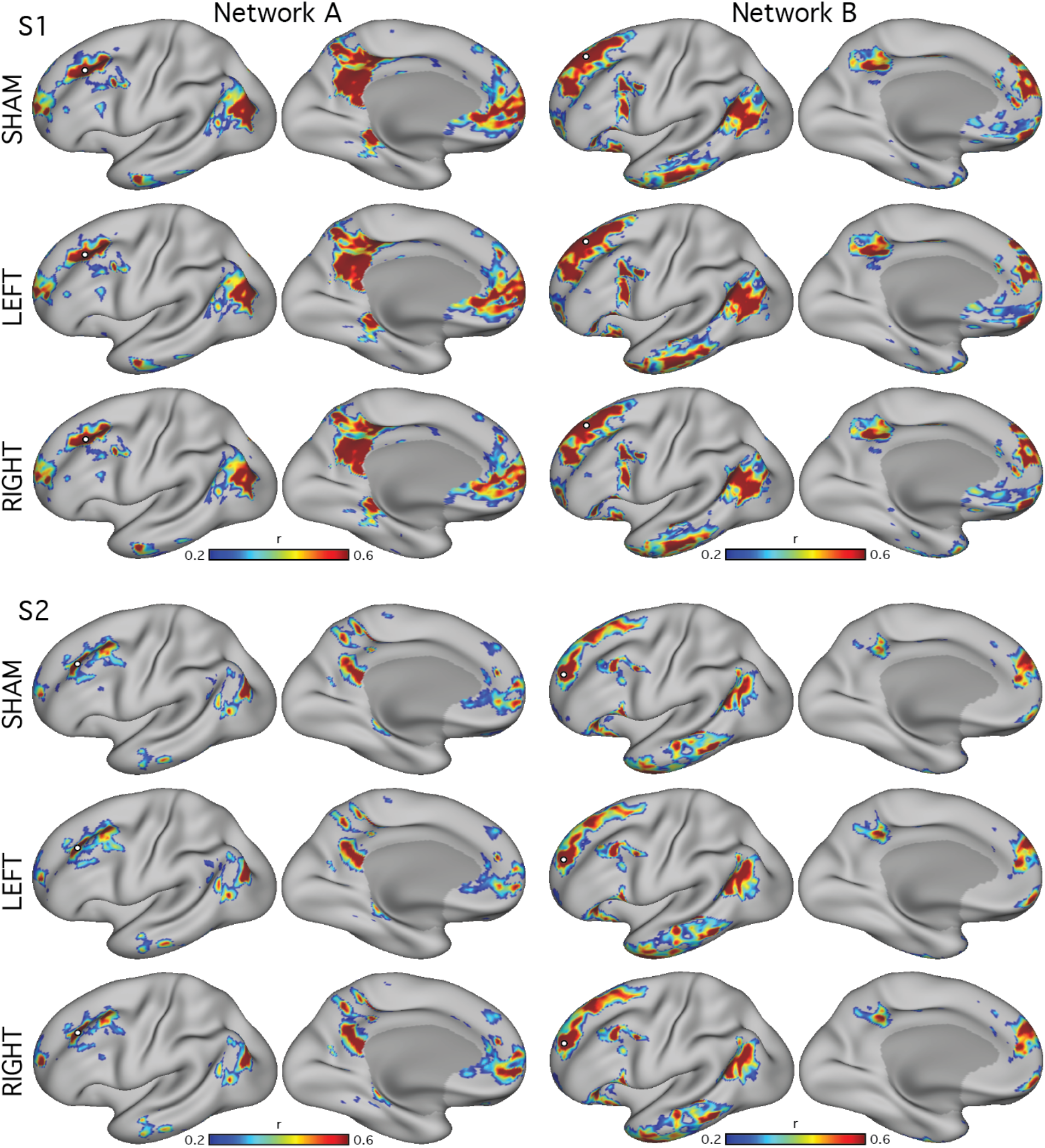
Negligible effects of transcranial magnetic stimulation (TMS) on functional connectivity. To confirm the decision to treat all TMS stimulation runs as standard rest fixation in the replication analyses (Fig. 1 and Supporting Figs. S1 and S2), functional connectivity maps were created using data grouped by stimulation condition, including sham (S1, n=19; S2, n=15), right(S1, n=19; S2, n=18), and left-hemisphere stimulation (S1, n=16; S2, n=17). Seed-based functional connectivity maps were created by averaging the maps within each of these conditions for S1 and S2. Similar maps were obtained for each condition, confirming that the detection of the two juxtaposed networks was not a consequence of the stimulation protocol.

## References

Amunts K, Schleicher A, Bürgel U, Mohlberg H, Uylings HB, Zilles K. Broca’s region revisited: Cytoarchitecture and intersubject variability. J Comp Neurol 412: 319–341, 1999.

Andersson JLR, Jenkinson M, Smith S. Non-linear registration, aka spatial normalisation. FMRIB technical report TR07JA2, 2010.

Andrews-Hanna JR, Saxe R, Yarkoni T. Contributions of episodic retrieval and mentalizing to autobiographical thought: Evidence from functional neuroimaging, resting-state connectivity, and fMRI meta-analyses. Neuroimage 91: 324–335, 2014.

Binder JR, Desai RH, Graves WW, Conant LL. Where is the semantic system? A critical review and metaanalysis of 120 functional neuroimaging studies. Cereb Cortex 19: 2767–2796, 2009.

Braga RM, Sharp DJ, Leeson C, Wise RJS, Leech R. Echoes of the brain within default mode, association and heteromodal cortices. J Neurosci 33: 14031–14039, 2013.

Braga RM, Buckner RL. Parallel interdigitated distributed networks within the individual estimated by intrinsic functional connectivity. Neuron 95: 457–471, 2017.

Brodmann K. Brodmann’s Localisation in the Cerebral Cortex: The Principles of Comparative Localisation in the Cerebral Cortex Based on Cytoarchitectonics (LJ Carey Trans.). New York: Springer, 1909/2006.

Buckner RL, Andrews-Hanna JR, Schacter DL. The brain’s default network: Anatomy, function, and relevance to disease. Ann N Y Acad Sci 1124: 1–38, 2008.

Buckner RL, Sepulcre J, Talukdar T, Krienen FM, Liu H, Hedden T, Andrews-Hanna JR, Sperling RA, Johnson KA. Cortical hubs revealed by intrinsic functional connectivity: Mapping, assessment of stability, and relation to Alzheimer’s disease. J Neurosci 29: 1860–1873, 2009.

Buckner RL, Margulies DS. Macroscale cortical organization and a default-like transmodal apex network in the marmoset monkey. BioRxiv doi: https://doi.org/10.1101/415141, 2018.

Burman KJ, Reser DH, Yu, HH, Rosa MGP. Cortical input to the frontal pole of the marmoset monkey. Cereb Cortex 21: 1712–1737, 2011.

Clark VP, Keil K, Maisog JM, Courtney S, Ungerleider LG, Haxby JV. Functional magnetic resonance imaging of human visual cortex during face matching: A comparison with positron emission tomography. Neuroimage 4: 1–15, 1996.

Córcoles-Parada M, Müller NCJ, Ubero M, Serrano-Del-Pueblo VM, Mansilla F, Marcos-Rabal P, Artacho-Pérula E, Dresler M, Insausti R, Fernández G, Muñoz-López M. Anatomical segmentation of the human medial prefrontal cortex. J Comp Neurol 525: 2376–2393, 2017.

Cox R. AFNI: software for analysis and visualization of functional magnetic resonance neuroimages. Comput Biomed Res 29: 162–173, 1996.

Cox R. AFNI. What a long strange trip it’s been. Neuroimage 62: 743–747, 2012.

Cox R, Saad Z. Surfing the Connectome: InstaCorr in AFNI and SUMA. International Conference on RestingState Functional Brain Connectivity, conference, 2010.

De Martino F, Esposito F, van de Moortele PF, Harel N, Formisano E, Goebel R, Ugurbil K, Yacoub E. Whole brain high-resolution functional imaging at ultra high magnetic fields: An application to the analysis of resting state networks. Neuroimage 57: 1031–44, 2011.

Doucet G, Naveau M, Petit L, Delcroix N, Zago L, Crivello F, Jobard G, Tzourio-Mazoyer N, Mazoyer B, Mellet E, Joliot M. Brain activity at rest: A multiscale hierarchical functional organization. J Neurophysiol 105: 2753–2763, 2011.

Duvernoy HM. The Human Brain. Surface, Three-dimensional Sectional Anatomy With MRI, and Blood Supply. 2nd Edition. New York: Springer, 1999.

Fedorenko E, Hsieh PJ, Nieto-Castañón A, Whitfield-Gabrieli S, Kanwisher N. New method for fMRI investigations of language: Defining ROIs functionally in individual subjects. J Neurophysiol 104: 1177–1194, 2010.

Fedorenko E, Duncan J, Kanwisher N. Language-selective and domain-general regions lie side by side within Broca’s area. Curr Biol 22: 2059–2062, 2012.

Fischl B, Sereno MI, Dale AM. Cortical surface-based analysis. II: Inflation, flattening, and a surface-based coordinate system. Neuroimage 9: 195–207, 1999.

Goldman-Rakic PS. Topography of cognition: Parallel distributed networks in primate association cortex. Annu Rev Neurosci 11: 137–156, 1988.

Gordon EM, Laumann TO, Gilmore AW, Newbold DJ, Greene DJ, Berg JJ, Ortega M, Hoyt-Drazen C, Gratton C, Sun H, Hampton JM, Coalson RS, Nguyen AL, McDermott KB, Shimony JS, Snyder AZ, Schlaggar BL, Petersen SE, Nelson SM, Dosenbach NUF. Precision functional mapping of individual human brains. Neuron 95: 791–807, 2017.

Hale JR, Brookes MJ, Hall EL, Zumer JM, Stevenson CM, Francis ST, Morris PG. Comparison of functional connectivity in default mode and sensorimotor networks at 3 and 7T. MAGMA 23: 339–349, 2010.

Henssen A, Zilles K, Palomero-Gallagher N, Schleicher A, Mohlberg H, Gerboga F, Eickhoff SB, Bludau S, Amunts K. Cytoarchitecture and probability maps of the human medial orbitofrontal cortex. Cortex 75: 87–112, 2016.

Hubel DH, Wiesel TN. Brain mechanisms of vision. Sci Am 241: 150–162, 1979.

Huth AG, de Heer WA, Griffiths TL, Theunissen FE, Gallant JL. Natural speech reveals the semantic maps that tile human cerebral cortex. Nature 532: 453–458, 2016.

Jenkinson M, Smith SM. A global optimisation method for robust affine registration of brain images. Med Image Anal 5: 143–156, 2001.

Jenkinson M, Bannister PR, Brady JM, Smith SM. Improved optimisation for the robust and accurate linear registration and motion correction of brain images. Neuroimage 17: 825–841, 2002.

Jenkinson M. Improving the registration of B0-disorted EPI images using calculated cost function weights. Tenth Int. Conf. on Functional Mapping of the Human Brain, 2004.

Kanwisher N. Functional specificity in the human brain: a window into the functional architecture of the mind. Proc Natl Acad Sci USA 107: 11163–11170, 2010.

Keil B, Triantafyllou C, Hamm M, Wald LL. Design optimization of a 32-channel head coil at 7T. In Proceedings of the 18th Annual Meeting of ISMRM, Montreal, Quebec. Abstract 1493, 2010.

Kong R, Li J, Orban C, Sabuncu MR, Liu H, Schaefer A, Sun N, Zuo XN, Holmes AJ, Eickhoff SB, Yeo BTT. Spatial topography of individual-specific cortical networks predicts human cognition, personality, and emotion. Cereb Cortex doi:10.1093/cercor/bhy123 [Epub ahead of print], 2018.

Kwong KK, Belliveau JW, Chesler DA, Goldberg IE, Weisskoff RM, Poncelet BP, Kennedy DN, Hoppel BE, Cohen MS, Turner R, Cheng HM, Brady T, Rosen BR. Dynamic magnetic resonance imaging of human brain activity during primary sensory stimulation. Proc Natl Acad Sci USA 89: 5675–5679, 1992.

Lafer-Sousa R, Conway BR. Parallel, multi-stage processing of colors, faces and shapes in macaque inferior temporal cortex. Nat Neurosci 16: 1870–1878, 2013.

Lafer-Sousa R, Conway BR, Kanwisher NG. Color-biased regions of the ventral visual pathway lie between face- and place-selective regions in humans, as in macaques. J Neurosci 36: 1682–1697, 2016.

Laumann TO, Gordon EM, Adeyemo B, Snyder AZ, Joo SJ, Chen MY, Gilmore AW, McDermott KB, Nelson SM, Dosenbach NU, Schlaggar BL, Mumford JA, Poldrack RA, Petersen SE. Functional system and areal organization of a highly sampled individual human brain. Neuron 87: 657–70, 2015.

Marcus DS, Harwell J, Olsen T, Hodge M, Glasser MF, Prior F, Jenkinson M, Laumann T, Curtiss SW, van Essen DC. Informatics and data mining tools and strategies for the human connectome project. Front Neuroinform 5: 4, 2011.

Margulies DS, Ghosh SS, Goulas A, Falkiewicz M, Huntenburg JM, Langs G, Bezgin G, Eickhoff SB, Castellanos FX, Petrides M, Jefferies E, Smallwood J. Situating the default-mode network along a principal gradient of macroscale cortical organization. Proc Natl Acad Sci USA 113: 12574–12579, 2016.

Mazziotta JC, Toga AW, Evans AC, Fox P, Lancaster J. A probabilistic atlas of the human brain: theory and rationale for its development. Neuroimage 2: 89–101, 1995.

Mennes M, Jenkinson M, Valabregue R, Buitelaar JK, Beckmann C, Smith S. Optimizing full-brain coverage in human brain MRI through population distributions of brain size. Neuroimage 98: 513–520R, 2014.

Michalka SW, Kong L, Rosen ML, Shinn-Cunningham BG, Somers DC. Short-term memory for space and time flexibly recruit complementary sensory-biased frontal lobe attention networks. Neuron 87: 882–892, 2015.

Moeller S, Freiwald WA, Tsao DY. Patches with links: A unified system for processing faces in the macaque temporal lobe. Science 320: 1355–1359, 2008.

Ogawa S, Tank DW, Menon R, Ellermann JM, Kim SG, Merkle H, Ugurbil K. Intrinsic signal changes accompanying sensory stimulation: Functional brain mapping with magnetic resonance imaging. Proc Natl Acad Sci USA 89: 5951–5955, 1992.

Ojemann JG, Akbudak E, Snyder AZ, mcKinstry RC, Raichle ME, Conturo TE. Anatomic localization and quantitative analysis of gradient refocused echo-planar fMRI susceptibility artifacts. Neuroimage 6: 156–167, 1997.

Öngür D, Price JL. The organization of networks within the orbital and medial prefrontal cortex of rats, monkeys and humans. Cereb Cortex 10: 206–219, 2000.

Öngür D, Ferry AT, Price JL. Architectonic subdivision of the human orbital and medial prefrontal cortex. J Comp Neurol 460: 425–449, 2003.

Peer M, Salomon R, Goldberg I, Blanke O, Arzy S. Brain system for mental orientation in space, time, and person. Proc Natl Acad Sci USA 112: 11072–11077, 2015.

Petrides M, Pandya DN. Comparative architectonic analysis of the human and macaque frontal cortex. In F. Boller, H. Spinnler (Eds) Handbook of Neuropsychology, Vol 9. Elsevier Science, Amsterdam, 1994.

Poldrack RA, Laumann TO, Koyejo O, Gregory B, Hover A, Chen MY, Gorgolewski KJ, Luci J, Joo SJ, Boyd RL, Hunicke-Smith S, Simpson ZB, Caven T, Sochat V, Shine JM, Gordon E, Snyder AZ, Adeyemo B, Petersen SE, Glahn DC, Reese Mckay D, Curran JE, Göring HH, Carless MA, Blangero J, Dougherty R, Leemans A, Handwerker DA, Frick L, Marcotte EM, Mumford JA. Long-term neural and physiological phenotyping of a single human. Nat Commun 6: 8885, 2015.

Polimeni JR, Bhat H, Witzel T, Benner T, Feiweier T, Inati SJ, Renvall V, Heberlein K, Wald LL. Reducing sensitivity losses due to respiration and motion in accelerated echo planar imaging by reordering the autocalibration data acquisition. Magn Reson Med 75: 665–679, 2016.

Power JD, Cohen AL, Nelson SM, Wig GS, Barnes KA, Church JA, Vogel AC, Laumann TO, Miezin FM, Schlaggar BL, Petersen SE. Functional network organization of the human brain. Neuron 72: 665–678, 2011.

Power JD, Schlaggar BL, Lessov-Schlaggar CN, Petersen SE. Evidence for hubs in human functional brain networks. Neuron 79: 798–813, 2013.

Raichle ME. The brain’s default mode network. Annu Rev Neurosci 38, 433–447, 2015.

Rajkowska G, Goldman-Rakic PS. Cytoarchitectonic definition of prefrontal areas in the normal human cortex: II. Variability in locations of areas 9 and 46 and relationship to the Talairach Coordinate System. Cereb Cortex 5: 323–337, 1995.

Rosa MGP, Soares JGM, Chaplin TA, Majka P, Bakola S, Phillips KA, Reser DH, Gattass R. Cortical afferents of area 10 in Cebus monkeys: Implications for the evolution of the frontal pole. Cereb Cortex Online: doi:10.1091/cercor/bhy044, 2018.

Rosene DL, Van Hoesen GW. Hippocampal efferents reach widespread areas of cerebral cortex and amygdala in the rhesus monkey. Science 198: 315–317, 1977.

Selemon LD, Goldman-Rakic PS. Common cortical and subcortical targets of the dorsolateral prefrontal and posterior parietal cortices in the rhesus monkey: Evidence for a distributed neural network subserving spatially guided behavior. J Neurosci 8: 4049–4068, 1988.

Setsompop K, Gagoski BA, Polimeni JR, Witzel T, Wedeen VJ, Wald LL. Blipped-controlled aliasing in parallel imaging for simultaneous multislice echo planar imaging with reduced g-factor penalty. Magn Reson Med 67: 1210–1224, 2012.

Smith SM, Jenkinson M, Woolrich MW, Beckmann CF, Behrens TEJ, Johansen-Berg H, Bannister PR, De Luca M, Drobnjak I, Flitney DE, Niazy R, Saunders J, Vickers J, Zhang Y, De Stefano N, Brady JM, Matthews PM. Advances in functional and structural MR image analysis and implementation as FSL. Neuroimage 23: 208–219, 2004.

Spiridon M, Fischl B, Kanwisher N. Location and spatial profile of category-specific regions in human extrastriate cortex. Hum Brain Mapp 27: 77–89, 2006.

Steinmetz H, Seitz, RJ. Functional anatomy of language processing: Neuroimaging and the problem of individual variability. Neuropsychologia 29: 1149–1161, 1991.

Tomasi D, Volkow ND. Functional connectivity density mapping. Proc Natl Acad Sci USA 107: 9885–9890, 2010.

van der Kouwe AJ, Benner T, Fischl B, Schmitt F, Salat DH, Harder M, Sorensen AG, Dale AM. On-line automatic slice positioning for brain MR imaging. Neuroimage 27: 222–230, 2005.

van der Kouwe AJ, Benner T, Salat DH, Fischl B. Brain morphometry with multiecho MPRAGE. Neuroimage 40: 559–569, 2008.

Vogt BA. Cingulate Neurobiology and Disease. Oxford: Oxford University Press, 2009.

Wang D, Buckner RL, Fox MD, Holt DJ, Holmes AJ, Stoecklein S, Langs G, Pan R, Qian T, Li K, Baker JT, Stufflebeam SM, Wang K, Wang X, Hong B, Liu H. Parcellating cortical functional networks in individuals. Nat Neurosci 18: 1853–1860, 2015.

Weiskopf N, Hutton C, Josephs O, Deichmann R. Optimal EPI parameters for reduction of susceptibility-induced BOLD sensitivity losses: A whole-brain analysis at 3 T and 1.5 T. Neuroimage 33: 493–504, 2006.

Yeo BTT, Krienen FM, Sepulcre J, Sabuncu MR, Lashkari D, Hollinshead M, Roffman JL, Smoller JW, Zöllei L, Polimeni JR, Fischl B, Liu H, Buckner RL. The organization of the human cerebral cortex estimated by intrinsic functional connectivity. J Neurophysiol 106: 1125–1165, 2011.

